# Neuroinvasion and anosmia are independent phenomena upon infection with SARS-CoV-2 and its variants

**DOI:** 10.1101/2022.08.31.505985

**Authors:** Guilherme Dias de Melo, Victoire Perraud, Flavio Alvarez, Alba Vieites-Prado, Seonhee Kim, Lauriane Kergoat, Anthony Coleon, Bettina Salome Trüeb, Magali Tichit, Aurèle Piazza, Agnès Thierry, David Hardy, Nicolas Wolff, Sandie Munier, Romain Koszul, Etienne Simon-Lorière, Volker Thiel, Marc Lecuit, Pierre-Marie Lledo, Nicolas Renier, Florence Larrous, Hervé Bourhy

**Affiliations:** Institut Pasteur, Université Paris Cité, Lyssavirus Epidemiology and Neuropathology Unit, F-75015, Paris, France; Institut Pasteur, Université Paris Cité, Channel Receptors Unit, F-75015 Paris, France; Sorbonne Université, Collège Doctoral, F-75005 Paris, France; Institut du Cerveau et de la Moelle Épinière, Laboratoire de Plasticité Structurale, Sorbonne Université, INSERM U1127, CNRS UMR7225, 75013 Paris, France; Institute of Virology and Immunology (IVI), Bern, Switzerland; Department of Infectious Diseases and Pathobiology, Vetsuisse Faculty, University of Bern, Bern, Switzerland; Institut Pasteur, Université Paris Cité, Histopathology Platform, F-75015 Paris, France; Institut Pasteur, Université Paris Cité, Spatial Regulation of Genomes Laboratory, F-75015 Paris, France; Institut Pasteur, Université Paris Cité, Molecular Genetics of RNA viruses Unit, F-75015 Paris, France; Institut Pasteur, Université Paris Cité, Evolutionary Genomics of RNA Viruses Group, F-75015 Paris, France; Multidisciplinary Center for Infectious Diseases, University of Bern, Bern, Switzerland; Institut Pasteur, Université Paris Cité, Inserm U1117, Biology of Infection Unit, 75015 Paris, France; Necker-Enfants Malades University Hospital, Division of Infectious Diseases and Tropical Medicine, APHP, Institut Imagine, 75006, Paris, France; Institut Pasteur, Université Paris Cité, Perception and Memory Unit, F-75015 Paris, France; CNRS, UMR3571, 75015 Paris, France

**Keywords:** host-pathogen interaction, host adaptation, COVID-19, neurotropism, olfaction, olfactory bulbs, central nervous system, reverse genetics

## Abstract

**SUMMARY:** Anosmia was identified as a hallmark of COVID-19 early in the pandemic, however, with the emergence of variants of concern, the clinical profile induced by SARS-CoV-2 infection has changed, with anosmia being less frequent. Here, we assessed the clinical, olfactory and neuroinflammatory conditions of golden hamsters infected with the original Wuhan SARS-CoV-2 strain, its isogenic ORF7-deletion mutant and three variants: Gamma, Delta, and Omicron/BA.1. We show that infected animals developed a variant-dependent clinical disease including anosmia, and that the ORF7 of SARS-CoV-2 contributes to the induction of olfactory dysfunction. Conversely, all SARS-CoV-2 variants were found to be neuroinvasive, regardless of the clinical presentation they induce. Taken together, this confirms that neuroinvasion and anosmia are independent phenomena upon SARS-CoV-2 infection. Using newly generated nanoluciferase-expressing SARS-CoV-2, we validated the olfactory pathway as a major entry point into the brain *in vivo* and demonstrated *in vitro* that SARS-CoV-2 travels retrogradely and anterogradely along axons in microfluidic neuron-epithelial networks.

**Graphical abstract:** 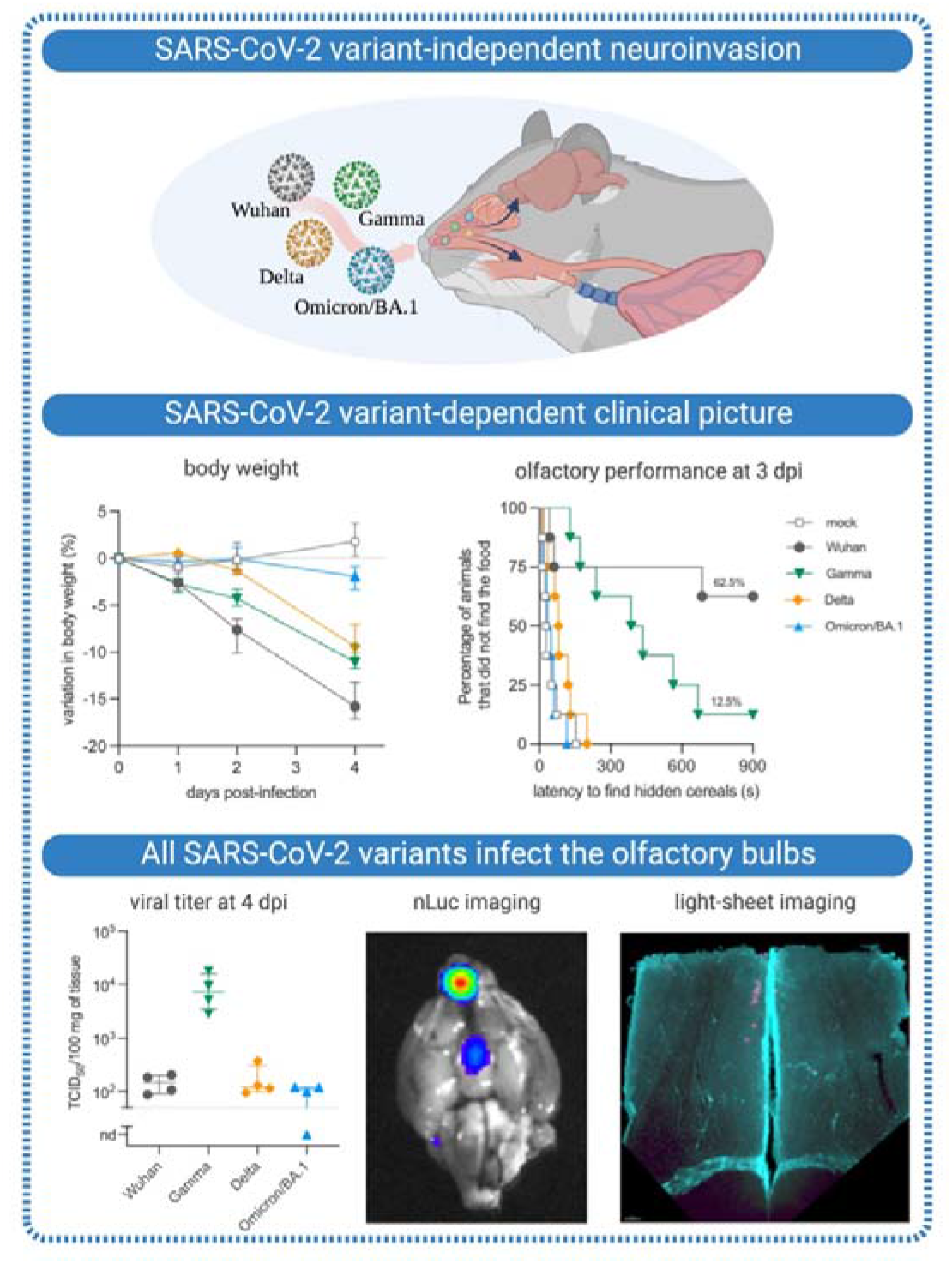

## INTRODUCTION

The COVID-19 pandemic remains a major global public health problem. Since the beginning of the pandemic in December 2019, more than 760 million cases have been confirmed, all SARS-CoV-2 variants combined ^1^. The original SARS-CoV-2 (Wuhan) gave rise to different variants of concern (VoCs) in the first year of the pandemic (Alpha/B.1.1.7, Beta/B.1.351, Gamma/P.1), which were almost totally replaced by the VoC Delta (B1.617.2, AY*) in 2020, by the VoC Omicron and its lineages (e.g. BA.1, BA.2 and BA.5) in 2021-2022 and recombinants (XBB*) ^1^.

SARS-CoV-2 infects cells of the upper and lower airways, and COVID-19 manifests by a multitude of respiratory and extra-respiratory symptoms, including neurological manifestations, ranging from headache and dizziness to anosmia, ageusia and even stroke ^2–4^. The neuropathology of COVID-19 is currently considered to be a consequence of inflammation and hypoxia, rather than direct viral invasion into the CNS ^5, 6^. However, with the emergence of the VoCs, and the increasing rate of vaccination or previous infection, the symptomatology of COVID-19 has changed. In this context, the clinical picture induced by the VoC Omicron/BA.1 is less severe or sometimes asymptomatic, with a higher rate of upper airways involvement, sparing the olfactory mucosa and consequently a lower incidence of anosmia, initially considered a hallmark of COVID-19 ^7–13^.

Previously, we determined that the infection of olfactory sensory neurons and loss of cilia in the olfactory mucosa caused by SARS-CoV-2 Wuhan in humans and Syrian hamsters were associated to olfaction loss, as well as to local inflammation and neuroinflammation in the olfactory bulbs, attesting that the golden hamster is a relevant model to study the pathogenesis of SARS-CoV-2 infection ^14–16^. Other authors reported persistent microgliosis in the olfactory bulbs of infected hamsters ^17^, and differences in the neuroinvasiveness and neurovirulence of VoCs have been described ^18^. Additionally, other authors have attempted to compare VoCs pathogenicity in hamsters ^19–21^, but studies relating SARS-CoV-2 brain invasion to neurological symptoms are lacking. In addition to the pathogenicity variation due to mutations in the spike of the variants, other viral proteins may play a role in anosmia and local inflammation, including ORF7, which has been shown to interfere with the host’s innate immunity ^22–24^, with cell adhesion in the olfactory mucosa ^25^, and with olfactory receptors ^26^.

Here, we identified that SARS-CoV-2 Wuhan and the VoCs Gamma, Delta and Omicron/BA.1 are all capable of invading the brain of Syrian hamsters and of eliciting a tissue-specific inflammatory response. Using reverse genetics-generated bioluminescent viruses, we demonstrate that SARS-CoV-2 infects the olfactory bulbs, but the clinical profile, including the olfactory performance, is highly dependent on the variant. Further, deletion of the ORF7ab sequence in the ancestral virus (Wuhan) reduces the incidence of olfaction loss without affecting the clinical picture nor the neuroinvasiviness via the olfactory bulbs. Accordingly, this work validates that SARS-CoV-2 may travel inside axons and the olfactory pathway as the main entry route by SARS-CoV-2 into the brain and corroborates the neurotropic potential of SARS-CoV-2 variants. Neuroinvasion and anosmia are therefore independent phenomena upon SARS-CoV-2 infection.

## RESULTS

### SARS-CoV-2 induces a clinical disease in hamsters, with a VoC-related severity difference

We first investigated the differences in the clinical picture induced by different SARS-CoV-2 VoCs in comparison with the ancestral virus (Wuhan). We first tested the *in vitro* growth curves of these viruses in Vero-E6 cells, and found no significant differences (Fig. S1A). Then, male golden hamsters were inoculated intranasally with 6x10^4^ PFU of SARS-CoV-2 Wuhan, or the VoCs Gamma, Delta or Omicron/BA.1 and followed them up for 4 days post-infection (dpi). All SARS-CoV-2- infected animals presented a progressive loss of weight; however, a variant effect was observed (Kruskal-Wallis P<0.0001, Fig. 1A-B), with SARS-CoV-2 Wuhan-infected animals presenting the most intense median weight loss (15.8%, interquartile range ‘IQR’ 3.9%), followed by Gamma-infected animals (11.0%, IQR 2.6%), Delta-infected animals (9.4%, IQR 3.1%) and Omicron/BA.1-infected animals (1.9%, IQR 2.5%). Non-specific sickness-related clinical signs (ruffled fur, slow movement, apathy) followed the same pattern (Kruskal-Wallis P<0.0001, Fig. 1D-E), with SARS-CoV-2 Wuhan infected animals presenting the worse clinical picture, followed by Gamma and Delta-infected animals, while Omicron/BA.1-infected animals presented a delayed manifestation of mild signs, clinically observable at 4 dpi only.

**Figure 1.**
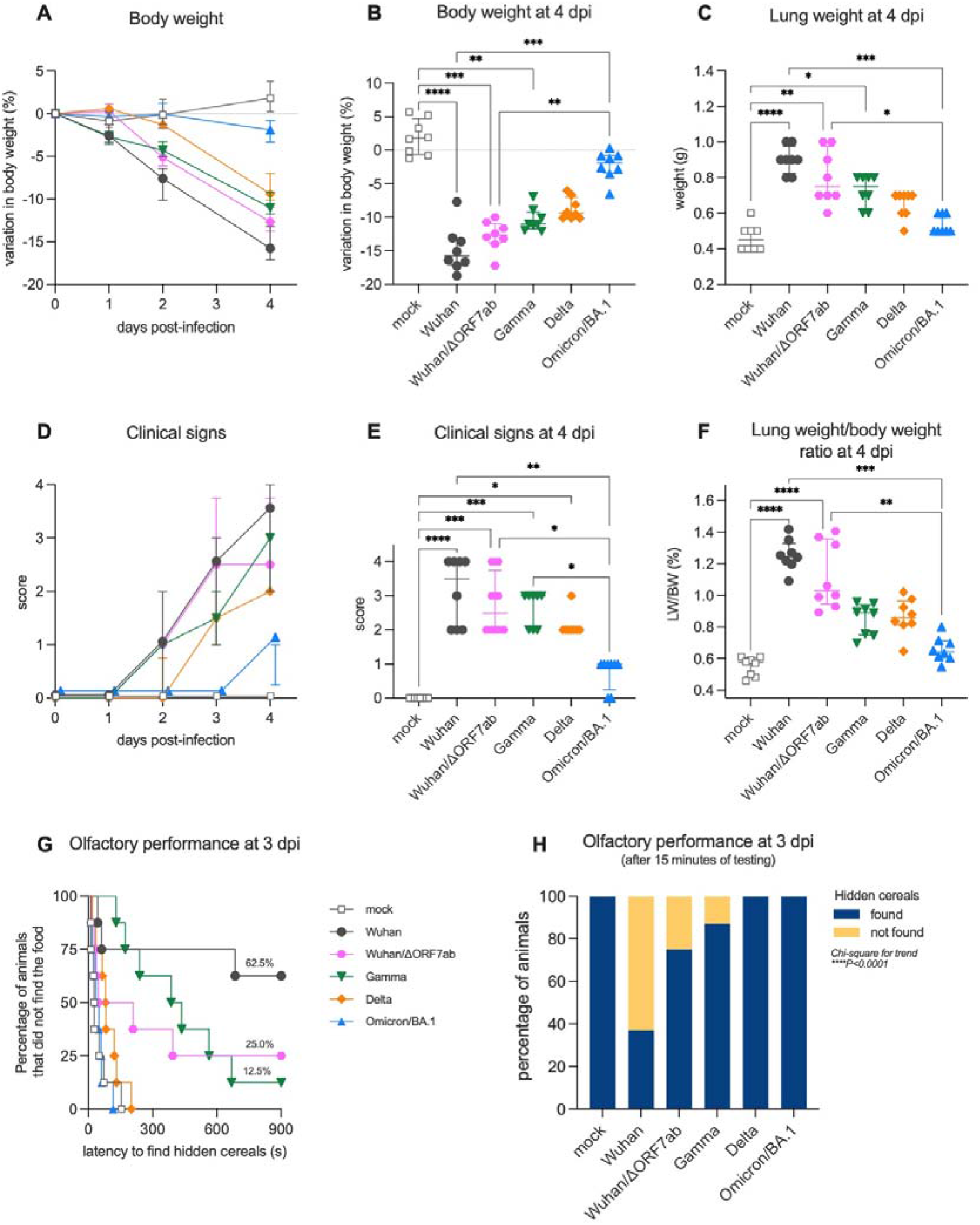
Clinical profile of hamsters infected with SARS-CoV-2 original virus (Wuhan), the recombinant Wuhan/ΔORF7ab or the variants of concern (VoC) Gamma, Delta, and Omicron/BA.1. **AB**. Body weight variation over four days post-infection. **C**. Lung weight measured at 4 dpi. **DE**. Clinical score over four days post-infection. The clinical score is based on a cumulative 0–4 scale: ruffled fur; slow movements; apathy; and absence of exploration activity. **F**. Lung weight to-body weight ratio measured at 4 dpi. Horizontal lines indicate median and the interquartile range (n=8/group). Kruskal-Wallis test followed by the Dunn’s multiple comparisons test (*P<0.05, **P<0.01, ***P<0.001, ****P<0.0001). **GH**. Olfactory performance measured at 3 days post infection (dpi). The olfaction test is based on the hidden (buried) food finding test. Curves represent the olfactory performance of animals during the test (G) and bars represent the final results (H) (n=8/group). Chi-square test for trend (****P<0.0001). See Supplementary Figures 1 and 2.

Olfaction loss is a typical clinical sign of SARS-CoV-2 Wuhan-infected hamsters ^14^, however olfactory deficit differed according to the different VoCs (Chi-square for trend P<0.0001): 62.5% (5/8) of SARS-CoV-2 Wuhan-infected animals presented loss of olfaction (Log-rank test compared to the mock P=0.0012), only 12.5% (1/8) of Gamma-infected animals lost olfaction completely with 62.5% (5/8) presenting an impaired olfactory performance (*i.e.* longer time to find the hidden cereals) (Log-rank test compared to the mock P<0.0001). In contrast, none of the Delta and Omicron/BA.1-infected animals presented signs of olfactory impairment (Fig. 1G-H). All animals found the visible food during the control test, demonstrating that no sickness behavior, visual impairment, or locomotor deficit was responsible for the delay in finding the hidden food.

Several mutations on the spike distinguish the different VoCs. Furthermore, some VoC isolates also present deletions in the ORF7 sequence ^27–30^, including the Delta isolate used in the present study (deletion spanning positions 27506 to 27540 of the reference SARS-CoV-2 Wuhan genome, which generates a stop codon; Fig. S1D). Since a possible link between ORF7b and olfaction loss has been proposed ^25^, we constructed a recombinant SARS-CoV-2 based on the CoV- 2/W backbone, where the ORF7ab sequence was replaced by that of GFP (SARS-CoV-2 Wuhan/ΔORF7ab) (Fig. S4A). Remarkably, the clinical profile exhibited by Wuhan/Δ ORF7ab infected hamsters was similar to the one caused by the wild-type SARS-CoV-2 Wuhan (Fig. 1A-F), showing that ORF7ab is not essential for viral infection and replication. However, the olfaction loss incidence decreased significantly; only 25% (2/8) of the infected animals presented signs of anosmia, in contrast to the 62.5% (5/8) observed in CoV-2/SARS-COV-2 Wuhan-infected hamsters (Fig. 1G-H). Despite this reduction in anosmia incidence, Wuhan/ΔORF7ab was still detected in the airways and in the olfactory bulbs of infected animals, with even higher titers (Fig. 2A-B).

**Figure 2.**
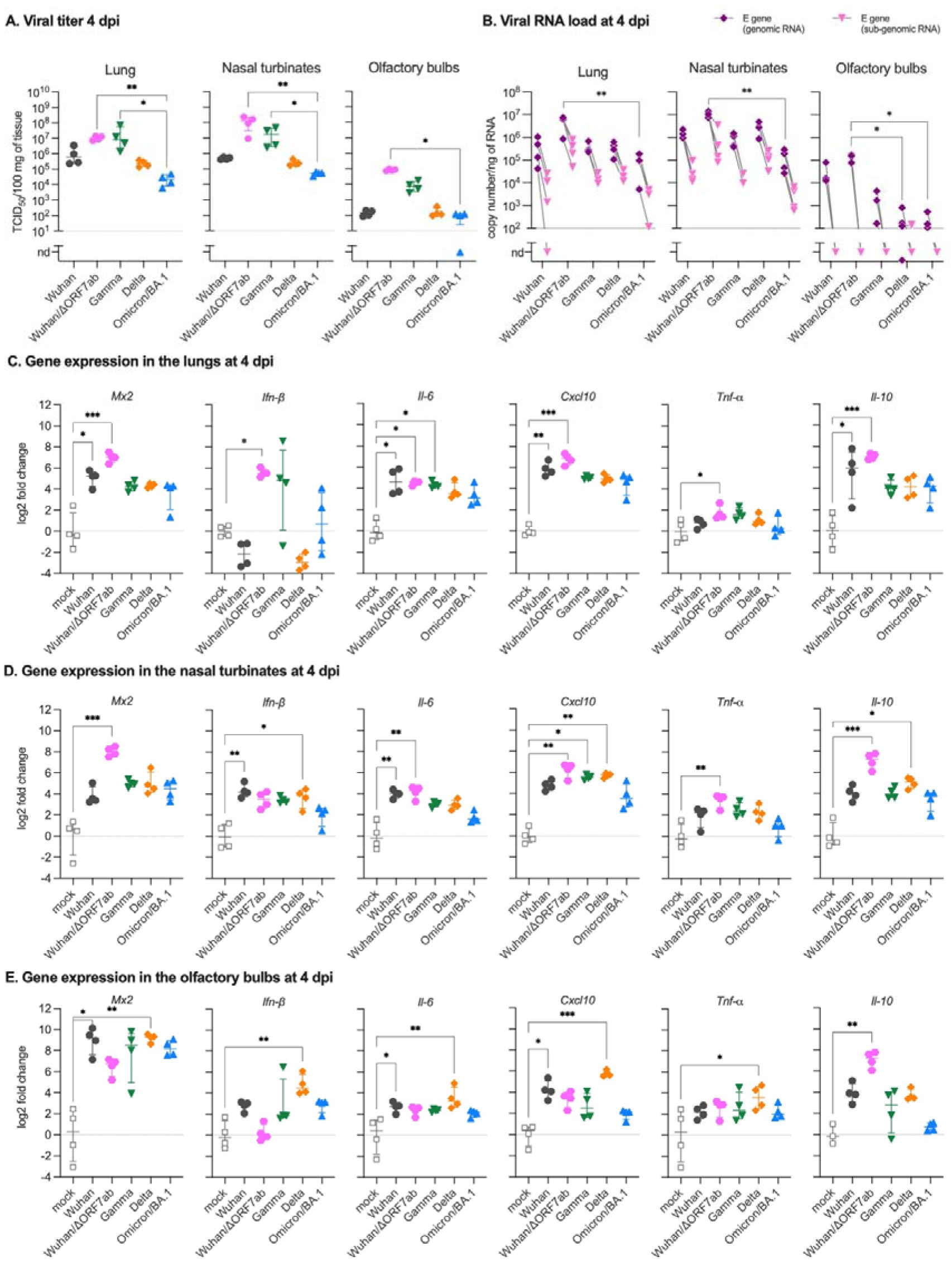
Virologic assessment and gene expression of selected immune mediators in different tissues of hamsters infected with SARS-CoV-2 original virus (Wuhan), the recombinant Wuhan/ΔORF7ab or the variants of concern (VoC) Gamma, Delta, and Omicron/BA.1. **A.** Infectious viral titers in the lung, nasal turbinates, and olfactory bulbs at 4 days post-infection (dpi) expressed as TCID_50_per 100 mg of tissue. Horizontal lines indicate median and the interquartile range (n=4/group). **B.** SARS-CoV-2 viral RNA load detected in the lung, nasal turbinates, and olfactory bulbs at 4 dpi. Genomic and sub-genomic viral RNA were assessed based on the E gene sequence. Horizontal lines indicate median and the interquartile range. Gray lines connect symbols from the same animals (n=4/group). **C-E.** Gene expression values in the lung (C), nasal turbinates (D) and olfactory bulbs (E) of *Mx2, Ifn-β, Il-6, Cxcl10, Tnf-α* and *Il-10* at 4 dpi. Horizontal lines indicate median and the interquartile range (n=4/group). Kruskal-Wallis test followed by the Dunn’s multiple comparisons test (*P<0.05, **P<0.01). nd: not detected. See Supplementary Figures 1 and 2.

Regarding lung pathology, as for the other clinical parameters, lung enlargement and lung weight-to-body weight (LW/BW) ratio were significantly higher in SARS-COV-2 Wuhan and in Wuhan/ΔORF7ab-infected animals; Gamma and Delta-infected animals presented intermediate values, whereas Omicron/BA.1-infected animals’ values were close to those of the ones of mock infected animals (Kruskal-Wallis P<0.0001, Fig. 1C and 1F). Likewise, the histopathological findings in the lungs followed the same tendency: congestion, edema, mononuclear cells infiltration, thickening of the alveolar walls and bronchiolar epithelium desquamation were observed in all infected animals (Fig. S2A), along with a diffuse SARS-CoV-2 nucleocapsid staining (Fig. S2B), with more severe alterations observed in the lungs of SARS-COV-2 Wuhan-infected animals than in Gamma-, Delta-, and Omicron/BA.1-infected hamsters.

In a multivariate analysis of clinical parameters recorded at 4 dpi, the two-first principal components explained 91.9% of sample variability. Body weight loaded negatively to PC1 (principal component) and negatively correlated with the clinical score and the LW/BW ratio, whereas the olfactory performance loaded positively to PC2 (Fig. S3A). In the principal component analysis (PCA) plot regarding tissue-related inflammation, the difference among groups was marked: mock and Omicron/BA.1-infected animals loaded homogeneously and relatively close, followed by the Delta infected animals. SARS-COV-2 Wuhan-, Wuhan/ΔORF7ab-, and Gamma-infected animals loaded in a more dispersed way, far away from the other VoCs, effect of the olfactory performance deficit and clinical severity (Fig. S3C).

### SARS-CoV-2-infected airways respond to the infection regardless of the viral variants

Next, we measured the viral titer and RNA loads in the upper airways (nasal turbinates), and in the lower airways (lung). Infectious viruses were detected in the nasal turbinates and in the lungs of all infected hamsters regardless of the VoC, however, a variant effect was observed, with the highest values for the Gamma-infected animals, and the lowest values for the Omicron/BA.1-infected animals (Kruskal-Wallis P<0.0001, Fig. 2A). Genomic and sub-genomic SARS-CoV-2 RNA were detected equally in the lungs of all infected animals; but differently in the nasal turbinates, regardless of the positivity of all samples, Delta-infected animals presented the highest viral load (Kruskal-Wallis P=0.0023, Fig. 2B).

Both the upper and lower airways responded to the infection by all VoCs (Fig. 2CD), but with a tissue-specific inflammatory signature: in the lungs, *Mx2, Il-6, Cxcl10* and *Il-10* were upregulated for all VoCs, and the highest in SARS-COV-2 Wuhan-infected animals (Fig. 2C). In the nasal turbinates, the gene expression of *Ifn-β* and *Il-6* presented the highest values in SARS-COV-2 Wuhan-infected animals and the lowest in Omicron/BA.1-infected animals. *Mx2*, *Cxcl10* and *Il-10* expression were the highest in Delta-infected animals while Omicron/BA.1-infected animals showed the lowest levels (Fig. 2D). Interestingly, Wuhan/ΔORF7ab infection induced a different lung inflammatory signature compared to Wuhan or the other VoCs, represented by a higher expression of *Mx2* and *Ifn-β* (Fig. 2C). In the nasal turbinates, Wuhan/ΔORF7ab infection also induced a higher expression of *Mx2,* but also *Il-6*, *Cxcl10, Tnf-α* and *Il-10* (Fig. 2D). To better appreciate the tissue *vs.* virus-related gene expression, we performed a multivariate analysis of these data. Interestingly, the two-first principal components explained 79.5% of sample variability (Fig. S3B). In the PCA plots, the nasal turbinates of all infected hamsters were loaded in proximity to each other (Fig. S3D), whereas the lungs were separated by an important effect of *Il-6, Cxcl10* and *Il-10* (Fig. S3D).

### SARS-CoV-2 neuroinvasion and neuroinflammation in the olfactory bulbs

After establishing the clinical and inflammatory profile of the infected animals, we aimed to assess the effect of infection on the olfactory bulbs. Remarkably, even if the olfactory performance differed according to the VoCs (Fig. 1G-H), positive viral titers were detected in the olfactory bulbs of animals from all infected groups, with Wuhan/ΔORF7ab and Gamma-infected animals presenting the highest titer at 4 dpi (Kruskal-Wallis P=0.0096, Fig. 2A). These findings were corroborated by the detection of genomic viral RNA in the olfactory bulbs of animals from all infected groups as well, but viral RNA load was higher in SARS-COV-2 Wuhan-infected animals (Kruskal-Wallis P=0.0104, Fig. 2B). Conversely, in these samples, the sub-genomic RNA was below the detection limit.

The olfactory bulbs presented an intrintriguing inflammatory profile. The antiviral *Mx2* gene, along with the inflammatory genes *Ifn-β*, *Il-6* and *Cxcl10* were highly upregulated in the olfactory bulbs of all infected hamsters (Fig. 2E), regardless of their olfactory performance in the food-finding test (Fig. 1 G-H), yet *Il-10* expression was highly upregulated in the olfactory bulbs of SARS-COV-2 Wuhan/ΔORF7ab-infected animals (compared to the mock). Unexpectedly, the gene expression of these selected targets tended to be higher in the olfactory bulbs of Delta-infected animals (Fig. 2E). In a multivariate analysis for tissue and virus-related gene expression, the olfactory bulbs from SARS-COV-2 Wuhan-infected animals tended to load in proximity to the corresponding nasal turbinates (effect of *Il-6*, *Cxcl10* and *Il-10*), whereas olfactory bulbs from Gamma-, Delta and Omicron/BA.1-infected animals loaded separately, possibly reflecting the impact of *Mx2*, *Tnf-α* and *Ifn-β* differential expression (Fig. S3D).

Having detected the virus in the olfactory bulbs using virologic and molecular techniques, we aimed to visualize the infection. We examined the SARS-CoV-2 Wuhan distribution in the whole brain by combining whole-head tissue clearing with light sheet microscopy imaging using iDISCO+ ^31^. At 4 dpi, all Wuhan-infected hamsters displayed a diffuse viral distribution in the nasal cavity: nasal turbinates and olfactory epithelium (Fig. 3A and Video S1). Along the same lines, all SARS- COV-2 Wuhan-infected animals presented SARS-CoV-2 in the olfactory bulbs in sparse neurons, localized in the proximity of the olfactory nerve entry point (Fig. 3B and Video S1). Using this technique, no infected cells were observed in other areas of the brain and no alteration in the vascular network was detected.

**Figure 3.**
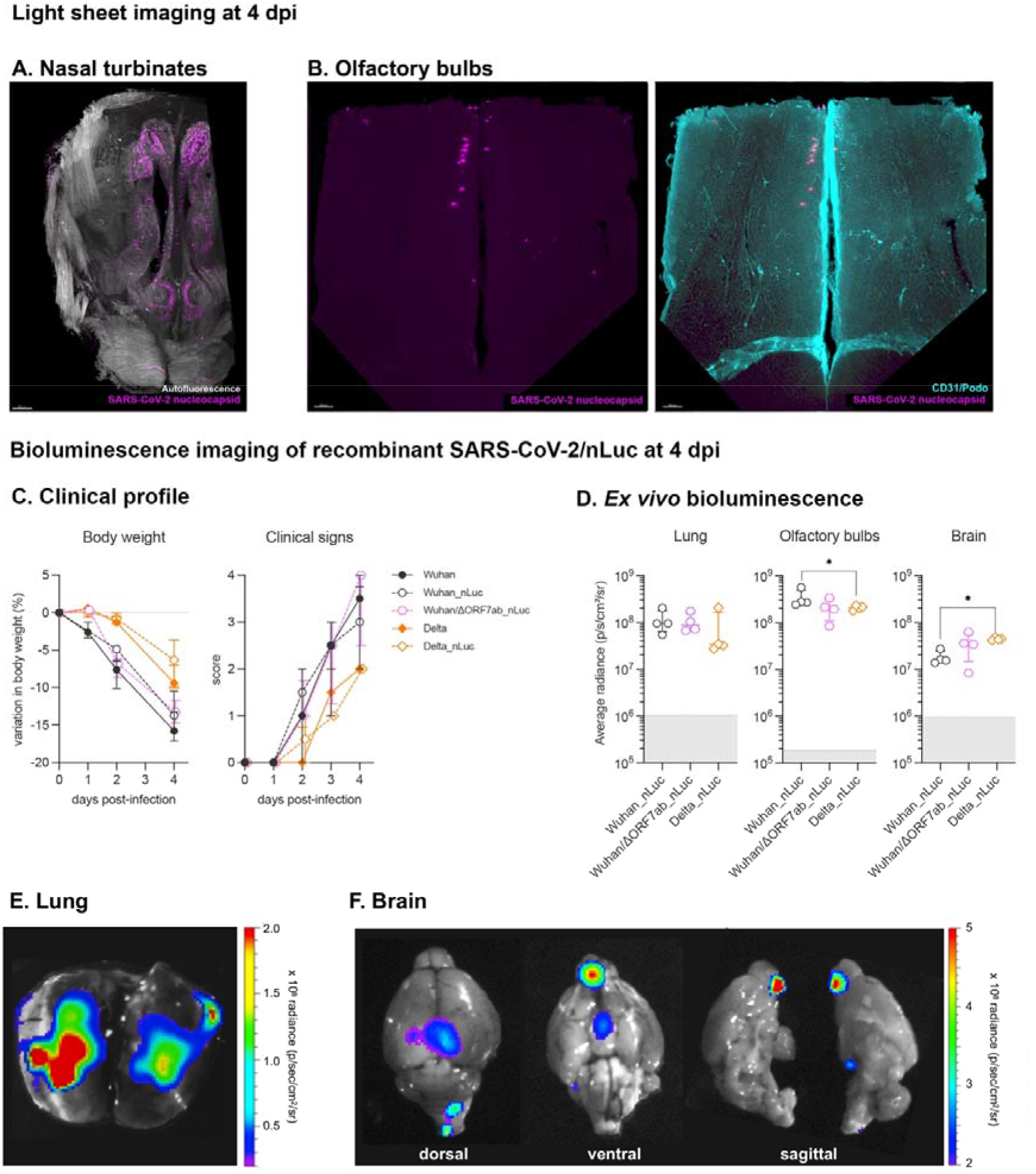
Imaging assessment of SARS-CoV-2 neuroinvasion. **AB.** iDISCO+ whole head clearing and immunolabeling against the SARS-CoV-2 nucleocapsid in hamsters infected with the SARS-CoV-2 original virus (Wuhan) at 4 days post-infection (dpi). **A.** Representative nasal turbinate (sagittal section from the skull) showing a diffuse distribution of SARS-CoV-2 (scale bar = 500 µm). **B.** Representative olfactory bulb section stained for SARS-CoV-2 nucleocapsid and for podoplanin (CD31/Podo) to identify the vascular compartment. Note presence of SARS-CoV-2 in olfactory bulb neurons, with no macroscopic alterations in the vascular network of the olfactory bulb. Scale bar: 200 µm. See supplementary video 1. **CF.** Bioluminescent recombinant SARS-CoV-2 viruses induce a clinical disease with infection of the olfactory bulbs. **C.** Clinical profile of hamsters infected with the recombinant viruses SARS-CoV-2 Wuhan_nLuc and Wuhan/ΔORF7ab_nLuc (based on the original SARS-CoV-2 Wuhan) and Delta_nLuc (based on the Delta variant). The body weight loss and the clinical score are comparable with those induced by wild-type viruses (n=4/group) (See Figure 1A). The clinical score is based on a cumulative 0–4 scale: ruffled fur; slow movements; apathy; and absence of exploration activity (See Figure 1D). **D.** *Ex vivo* bioluminescence values from the lungs, the olfactory bulbs (ventral view), the brains (ventral view) at 4 dpi (n=4/group). Horizontal lines indicate median and the interquartile range. Kruskal-Wallis test followed by the Dunn’s multiple comparisons test (*P<0.05). The gray crosshatched area corresponds to background signals obtained from the same tissues of a mock-infected hamster. **EF.** Representative *ex vivo* imaging of a lung (C) and a brain (D) of a hamster infected with a recombinant SARS-CoV-2 expressing the nLuc at 4 dpi. Note that in the brain, the major bioluminescent focus is localized in the ventral face of olfactory bulbs. See Supplementary Figure 3.

We next used a complementary method to visualize the infection of the olfactory bulbs. To this end, we generated recombinant viruses that express the nanoluciferase by reverse genetics (Wuhan_nLuc, Wuhan/ΔORF7ab_nLuc, and Delta_nLuc; Fig. S1B-C and S4A). The disease profile induced by these three novel recombinant viruses was similar to the profile induced by the wild- type parental viruses (Fig. 3C). *In vivo* imaging of infected hamsters was performed at 4 dpi using fluorofurimazine (FFz) as substrate. Very low positive signals were observed in the nasal cavity region, possibly due to the thickness of the skin and bones that could block the light emission, but significantly higher in the Wuhan_nLuc than in the Wuhan/ΔORF7ab_nLuc the Delta_nLuc (Kruskal Wallis P=0.0107, Fig. S4BC). More sensitive results were obtained during the *ex vivo* imaging. After euthanasia, the lungs and brains were collected and imaged. Positive signals in the lungs were obtained at the same intensity for the three viruses (Fig. 3D), and presented a multi-focal/diffuse distribution (Fig. 3E and S4D). The brains were imaged in the ventral position in order to better expose the olfactory bulbs. Intense luminescent signals were recorded from the brains of hamsters infected with all viruses; when placing the ROI (region of interest) exclusively in the olfactory bulbs, intensity was higher in Wuhan_nLuc compared to Delta_nLuc; conversely, when considering the whole brain ventral area, intensity was higher in Delta_nLuc. Wuhan/ΔORF7ab_nLuc induced intermediate values (Fig. 3C and S4D), which might reflect the higher replication rate detected in the olfactory bulbs infected Wuhan (Fig. 2B). Despite intensity differences, the olfactory bulbs were the major infected structure in the brain, and sagittal views of the brain indicate that the origin of the luminescent signal is the ventral part of the olfactory bulbs, which is situated above the olfactory epithelium in the nasal cavity (Fig. 3E and S4E).

### Anosmia is not associated with viral load or infection of the olfactory bulbs

All tested SARS-CoV-2 variants were found to be able to invade the CNS and infect the olfactory bulbs, but the incidence of olfactory dysfunctions varied according to the VoC, with hamsters infected with Delta and Omicron/BA.1 presenting no signs of anosmia (Fig. 1G-H). To test if the viral inoculum could influence the neuroinvasiveness of SARS-CoV-2 and the incidence of olfactory dysfunctions, we infected hamsters with a 2-log lower infectious dose of Wuhan (6x10^2^ PFU). These animals were subjected to the same clinical-behavioral tests described above. In the acute phase of the infection, up to 4 dpi, the low infectious dose induced similar body weight loss and clinical score as the animals infected with the initial inoculum, as well as infectious viral titers in the airways and in the olfactory bulbs (Fig. S5). Surprisingly, despite the presence of infectious virus in the olfactory bulbs, animals infected with a lower infectious dose presented a lower incidence of olfactory dysfunction (25%, 2/8).

### SARS-CoV-2 travels retrogradely and anterogradely along axons

*In vitro*, monocultures of neurons are hardly infectable by SARS-CoV-2 and co-culture with permissive cells is necessary ^32^. Consequently, we developed a co-culture system with hNSC-derived neurons and SARS-CoV-2-receptive epithelial cells (A549-hACE2-TMPRSS2 cells). We grew these co-cultures using axonal diodes in microfluidic devices ^33^ to assess the ability of SARS-CoV-2 to infect neurons and to move inside axons. In these devices, the co-cultures form neuron-epithelial networks in both the left and the right chambers of the device (Fig. 4A-B), that are connected exclusively via an unidirectional growth of axons from the left towards the right chamber (Fig. 4C). Then, the right or the left chamber of the microfluidic devices was infected with the ancestral virus (Wuhan), the Wuhan/ΔORF7ab and the VoCs Gamma, Delta and Omicron/BA.1. At 3 dpi, infected cells were detected in both the right and the left chambers of all devices, attesting that the virus travelled from the right to the left chamber inside the axons (retrograde axonal transport; Fig. 4D-H and S6), and from the left to the right chamber inside the axons (anterograde axonal transport; Fig. 4I-M and S7). To further confirm the involvement of the axonal transportation of SARS-CoV-2, we used ciliobrevin D, an inhibitor of the cytoplasmic dynein transport machinery which acts on the bidirectional transport of organelles along axons ^34^. When the infection occurred in the presence of ciliobrevin D, the retrograde transport was blocked, and no SARS-CoV-2-infected cell was observed in the left chamber (Fig. S8). To quantify the arrival of viral particles via the axons, we infected microfluidic devices in both retrograde and anterograde conditions with the recombinant viruses expressing nanoluciferase Wuhan_nLuc, Wuhan/ΔORF7ab_nLuc, and Delta_nLuc, and we measured the luminescence in the supernatants on a daily basis (Figure S9). Positive nLuc activity was detected in the supernatants of all receiving chambers from 1 dpi, increasing progressively overtime (Figure S9G and S9H), indicative of retrograde and anterograde SARS-CoV-2 movement. The luminescence values in the infected chambers were constant and at the maximum limit of detection (Fig. S1).

**Figure 4.**
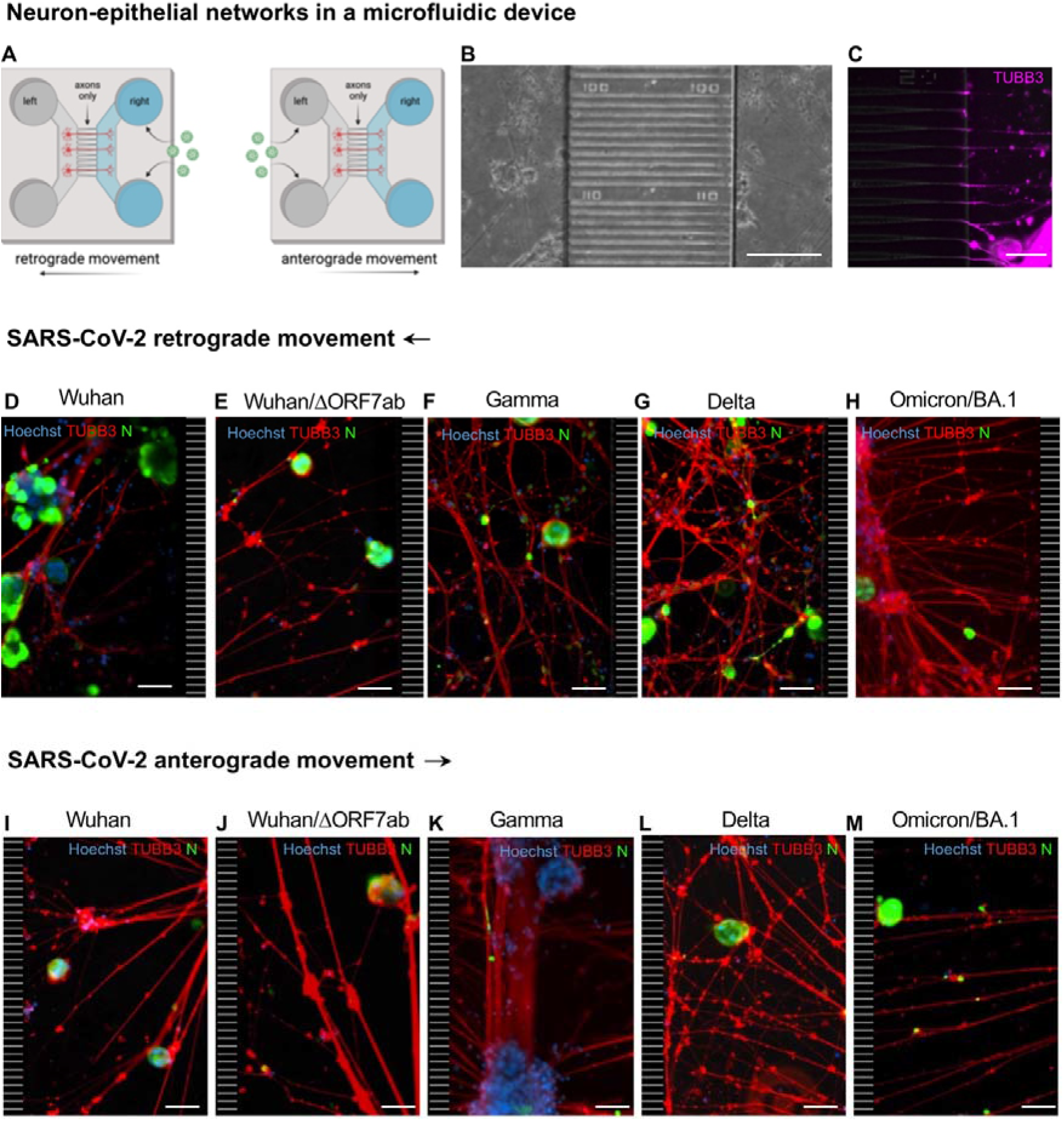
SARS-CoV-2 travels inside axons in *in vitro* neuron-epithelial networks. **A.** Schematic view of an axon diode in a microfluidic device where the left and the right chambers are connected exclusively by axons. In this study, SARS-CoV-2 was added in the right chamber (to assess the retrograde axonal transport) or in the left chamber (to assess the anterograde axonal transport). **B.** Bright field view of a neuron-epithelial network showing neuronal and A549-ACE2-TMPRSS2 co-culture in both chambers, separated by funnel-shaped micro-channels (scale bar = 200 µm). **C.** Detail of β-Tubulin III-positive axons crossing the funnel-shaped micro-channels and entering in the right chamber (scale bar = 50 µm). **D-H.** Neuron-epithelial networks infected at the right chamber with SARS-CoV-2 Wuhan (D), Wuhan/ΔORF7ab (E), Gamma (F), Delta (G) and Omicron/BA.1 (H) showing infected cells in the left chambers (scale bar = 50 µm). **I-M.** Neuron-epithelial networks infected at the left chamber with SARS-CoV-2 Wuhan (I), Wuhan/ΔORF7ab (J), Gamma (K), Delta (L), and Omicron/BA.1 (M) showing infected cells in the right chambers (scale bar = 50 µm). Hoechst: nuclei (blue). TUBB3: β-Tubulin III (red). N: SARS-CoV-2 nucleocapsid (green). The black and white striped zones in the right (D-H) and in the left (I-M) of the photomicrographs represents the microchannels, which are not accessible during immunostainings. See Supplementary Figures 6- 9.

## DISCUSSION AND CONCLUSIONS

The neurotropism of SARS-CoV-2 is still a matter of debate ^35^. While studies on natural infection in humans and experimental infection in animal models reported brain infection by SARS-CoV-2 ^14, 36–40^, including in *in vitro* human models of infection ^41–43^, others failed to detect SARS-CoV-2 in the nervous tissue ^5, 44^. This inconsistency may be due to the time of infection, as in most cases the available samples are post-mortal, and we have demonstrated, in the hamster model, that viral isolation is time-dependent, with higher viral loads found in acute time-points^14^. Despite this open question, the impact of SARS-CoV-2 infection on the brain is undeniable ^2, 45–47^. Most of these published data is related to the original SARS-CoV-2 (Wuhan), however, less information concerning VoCs neuropathogenesis is available ^10^. Here we show that all evaluated SARS-CoV-2 viruses (Wuhan, Gamma, Delta and Omicron/BA.1), including novel generated recombinant SARS-CoV-2, are able to invade the brain, most likely via the olfactory nerves, and of promoting inflammation in the nervous tissue.

Despite this shared neuroinvasiveness via the olfactory bulbs, disease profile and airways responses are quite dependent on the SARS-CoV-2 variant. Indeed, we show that all VoCs can infect golden hamsters and promote lung inflammation, including Omicron/BA.1 differently from other reports ^19, 48^, probably due to infectious doses or viral isolates. A “variant-effect” in the clinical presentation and in the tissue-related inflammation was evident, with disease severity presenting the following order: SARS-CoV-2 Wuhan > Gamma > Delta > OmicronBA.1, which supports the hypothesis that Omicron/BA.1 evolution has resulted in a tropism more restricted to the upper respiratory tract, thereby causing a less severe clinical disease ^10, 49–51^. Although with different severities, neuroinvasiveness, neurotropism and neurovirulence ^35^ seem to be conserved features among SARS-CoV-2 variants. Indeed, the data presented herein give support to the neuroinvasive ability of SARS-CoV-2, as besides detection of viral RNA and isolation of infectious virus from the olfactory bulbs, we could clearly observe SARS-CoV-2 infected neurons in the brain of hamsters by light sheet microscopy. Nevertheless in hamsters, unlike what has been reported in K18-hACE2 mice which ACE2 expression pattern is non-physiological and ectopic ^37^, detection of SARS-CoV-2- infected cells was restricted to the olfactory bulbs, without evidence of brain vasculature remodeling nor virus associated with blood vessels.

Olfactory bulb infection is therefore a common feature in the SARS-CoV-2 infectious process, regardless of the variant considered. An inflammation response was also observed in this tissue, with a common upregulation of the antiviral gene *Mx2*, regardless of the VoC. The reason why some VoCs do not cause olfaction loss is still an open question. The infection and inflammation of the olfactory bulbs, if involved in SARS-CoV-2-associated anosmia, does not seem to be enough to cause olfaction loss in the golden hamster. Aggressions to the olfactory mucosa, rather than the olfactory bulb, are indeed likely the main factors contributing to anosmia, as recovery from anosmia has been related to regeneration of the olfactory epithelium in hamsters ^52, 53^. Moreover, significant changes in the olfactory epithelium, such as apoptosis, architectural damages, inflammation, downregulation of odorant receptors and functional changes in olfactory sensory neurons, might also contribute, on top of the infection *per se*, to the occurrence of anosmia ^10, 14, 54– 57^. Therefore, it seems that infection and inflammation combined are needed to trigger anosmia, and the lower incidende of anosmia observed after infection by some VoCs is therefore linked to lower levels of inflammation.

Furthermore, besides mutations in the spike sequence in the SARS-CoV-2 variants genome, additional mutations may be present in other regions of the viral genome, including deletions in the ORF7a and ORF7b sequences ^27–30^. ORF7a and ORF7b have been related to viral-induced apoptosis, and to interference with the innate immunity and the antiviral response ^22–24, 58, 59^, without, however, being essential to viral infection and replication ^60, 61^. Further, ORF7b has the potential to interfere with cell adhesion proteins in the olfactory mucosa ^25^, and interestingly, a binary interaction between ORF7b and the human olfactory receptor OR1D5 has also been reported ^26^. Deletions or mutations in these regions may therefore play an additional role in the induction of olfactory disturbances, by preserving the architecture and structures of the olfactory epithelium. Another important point is that we can observe a similar clinical profile and low incidence of olfactory disturbances as SARS-CoV-2 Wuhan/ΔORF7ab by simply reducing the viral inoculum of SARS-CoV-2 Wuhan. Regardless of a comparative evolution of body weight loss and clinical score ^62^, anosmia was less frequent in animals infected with a low dose of SARS-CoV-2 Wuhan. This might be related to the less severe initial aggression suffered by the olfactory epithelium due to lower infectious doses ^53, 63^ and may also explain why other studies did not detect olfaction dysfunction despite detecting inflammatory changes in the brain ^64^.

Despite the occurrence or not of anosmia, SARS-CoV-2 can infect olfactory sensory neurons and the olfactory bulb ^10, 14^, and we show herein that the virus can infect neurons and travel inside the axons in both retrograde and anterograde directions. SARS-CoV-2 neuroinvasion was hypothesized to occur by axonal transport via cranial nerves (olfactory, vagus, trigeminal) or by the hematogeneous route ^35, 65, 66^. The data presented herein do not support hematogenous diffusion of SARS-CoV-2 towards the brain, but corroborate the hypothesis of the olfactory pathway as a preferential portal of entry towards the olfactory bulbs ^67^. Brain infection via the olfactory pathway therefore seems a common feature of coronaviruses ^68, 69^, regardless of clinical disease presentations. Finally, this study highlights that neuroinvasion and anosmia are therefore independent phenomena upon SARS-CoV-2 infection.

---

**Supplementary Figure 1.**
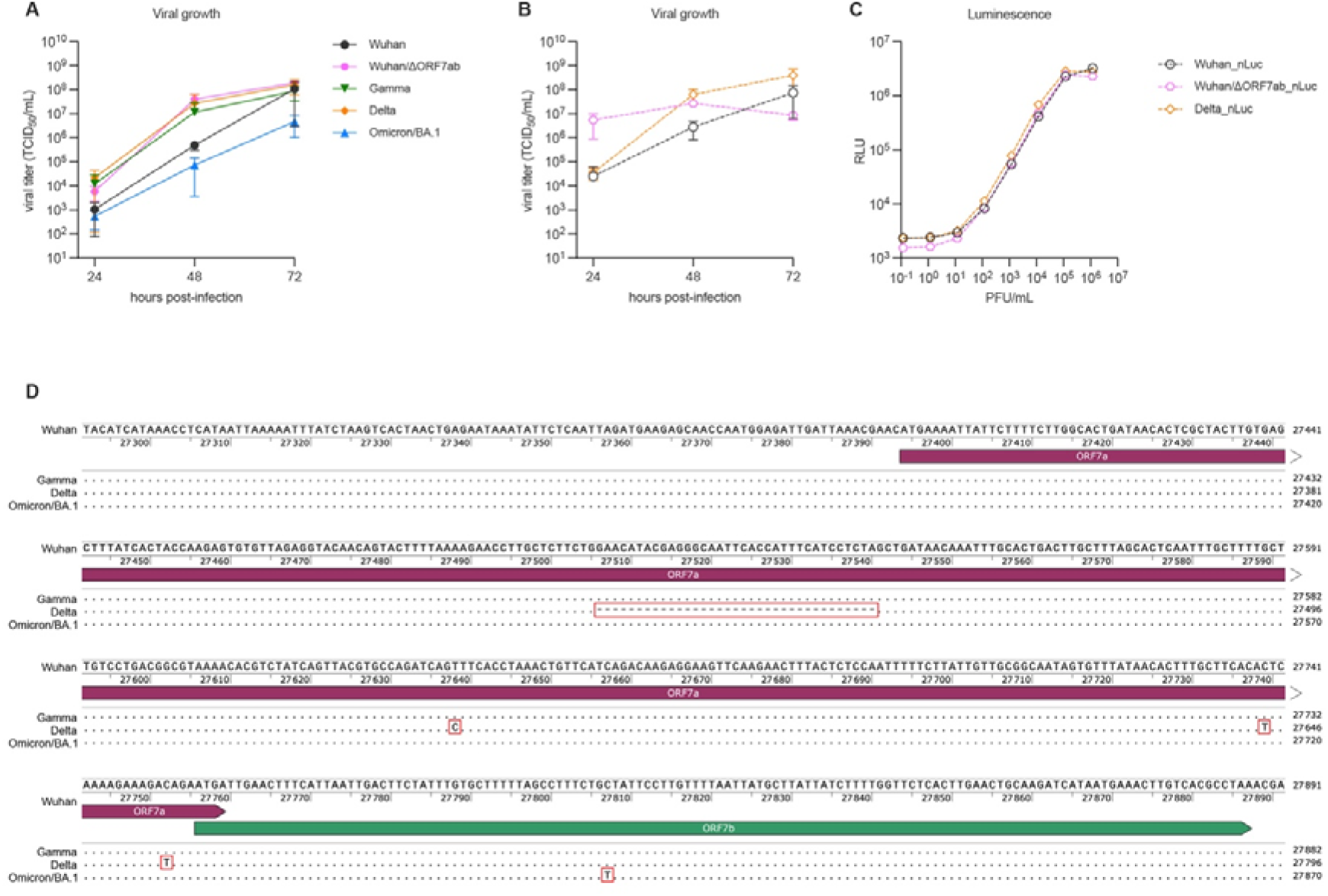
*In vitro* viral growth curves of different SARS-CoV-2 viruses. **A.** Growth curves of the SARS-CoV-2 original virus (Wuhan), the recombinant Wuhan/ΔORF7ab, or the variants of concern (VoC) Gamma, Delta and Omicron/BA.1. **B.** Growth curves of the recombinant nanoluciferase-expressing SARS-CoV-2 viruses Wuhan_nLuc, Wuhan/ΔORF7ab_nLuc, and Delta_nLuc. Horizontal lines indicate median and the interquartile range (n=3 independent replicates/time-point). Viral titers are expressed as the median tissue culture infectious dose (TCID50)/mL. **C**. Dilution curves showing the luminescence activity of the recombinant nanoluciferase-expressing SARS-CoV-2 viruses Wuhan_nLuc, Wuhan/ΔORF7ab_nLuc, and Delta_nLuc. RLU: relative light units. **D**. Multiple sequence alignment of the ORF7ab regions of Gamma, Delta and Omicron/BA.1 in comparison with the SARS-CoV-2 original virus (Wuhan). Deletions and mutations are indicated inside red squares. Related to Figures 1, 3, and 4.

**Supplementary Figure 2.**
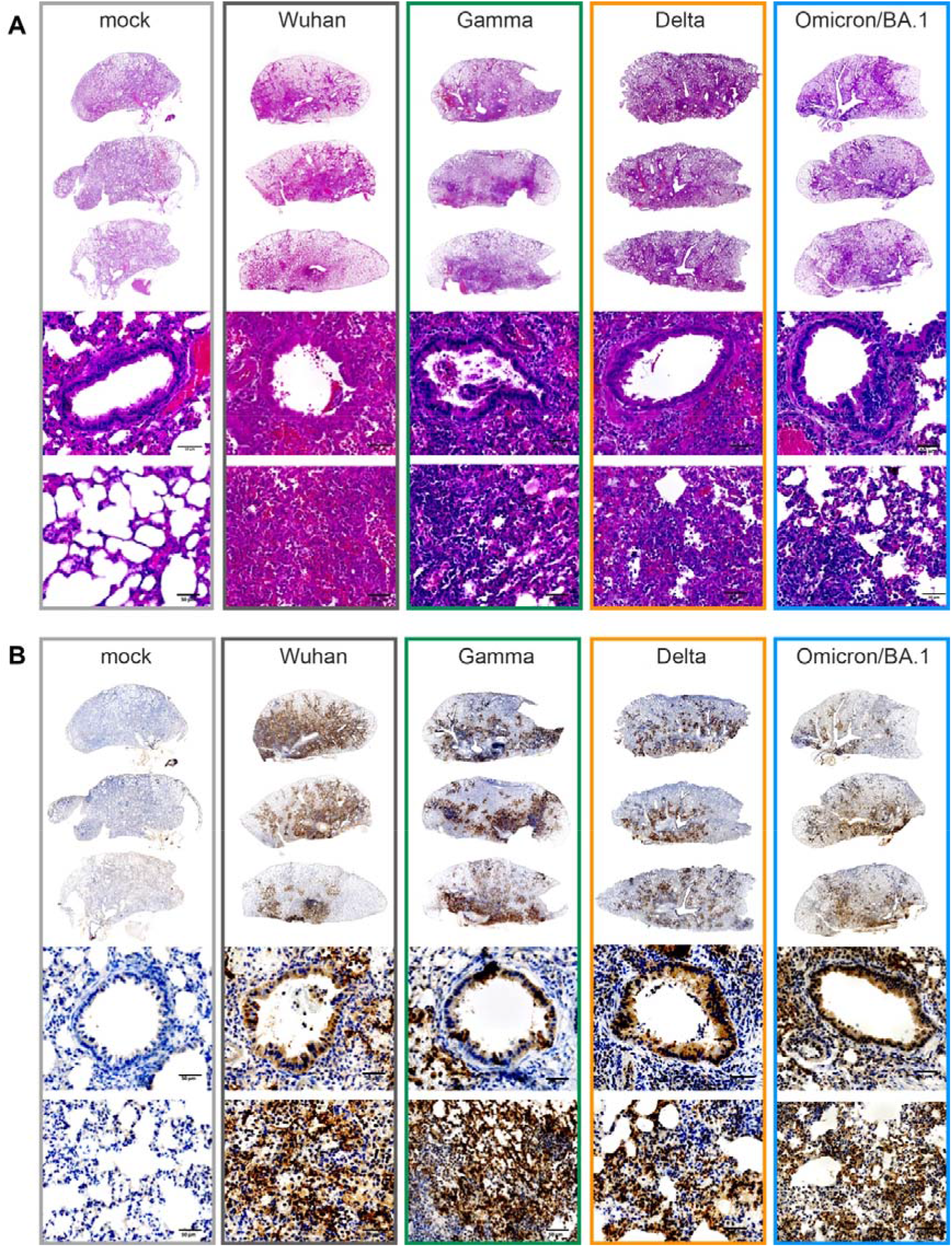
Histopathology and immunohistochemical aspects of freshly-collected lungs from hamsters infected with SARS-CoV-2 original virus (Wuhan) or the variants of concrn (VoC) Gamma, Delta and Omicron/BA.1 at 4 days post-infection (dpi). **A.** Representative images of Hematoxylin and Eosin (H&E) stained-whole lung sections (upper panels), bronchiolar epithelium (middle panels) and alveoli (bottom panels). **B.** Representative images of whole lung sections (upper panels), bronchiolar epithelium (middle panels) and alveoli (bottom panels) immuno-stained with SARS-CoV-2 N antibody. Immunoperoxidase, scale bar = 50 μm (n=8/group, except Omicron/BA.1 where n=4). Related to Figure 1.

**Supplementary Figure 3.**
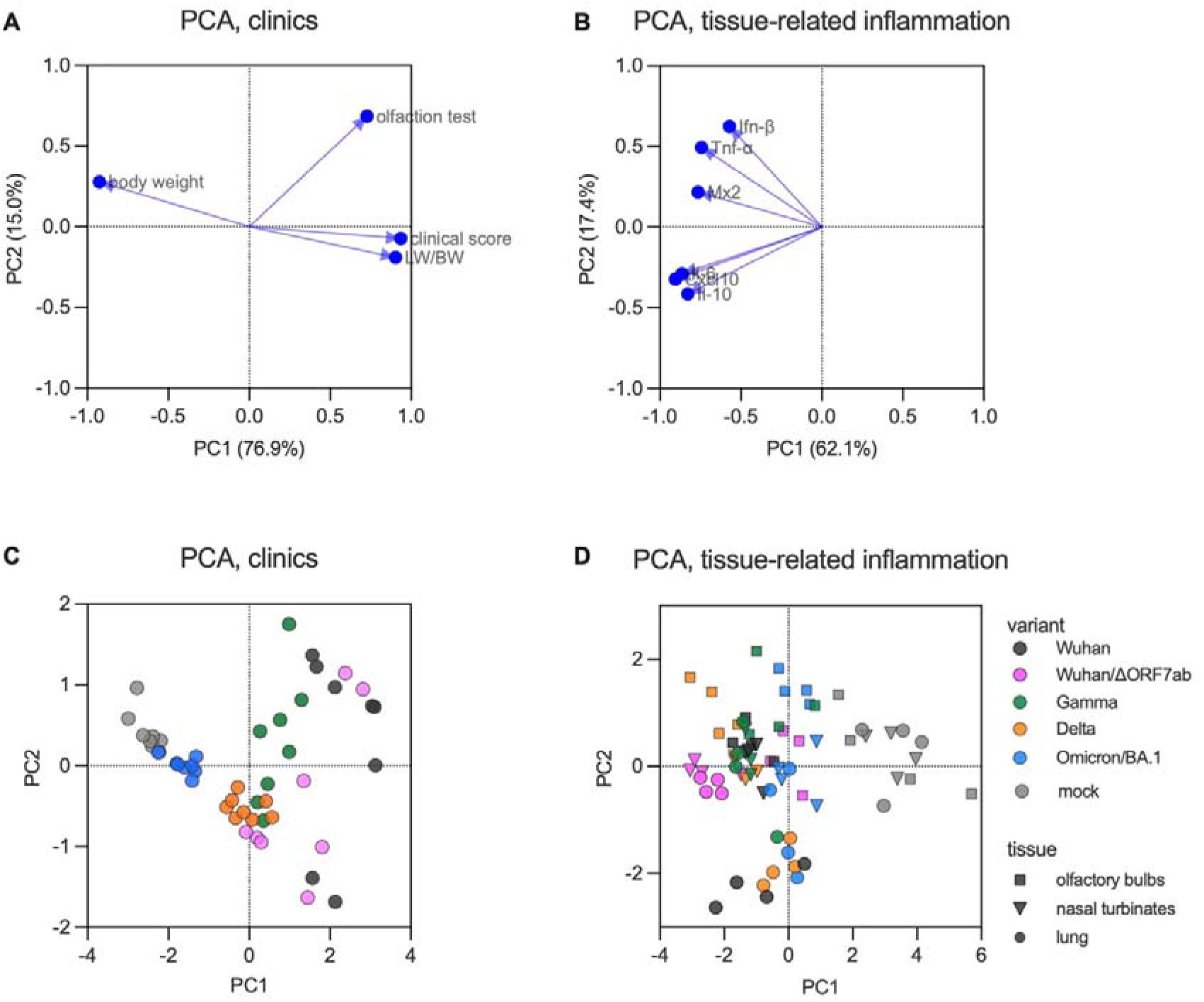
Principal component analysis (PCA) of clinical parameters and immune mediators in different tissues of hamsters infected with SARS-CoV-2 original virus (Wuhan), the recombinant Wuhan/ΔORF7ab or the variants of concern (VoC) Gamma, Delta and Omicron/BA.1. **A.** Variable correlation plot showing the correlation of clinical parameters in hamsters at 4 days post-infection (body weight variation, clinical score, olfaction test and lung weight-to-body weight ratio ‘LW/BW’). The two-first principal components explained 92.2% of sample variability (n=8/group). **B.** Variable correlation plot showing the correlation of immune mediators’ gene expression (*Mx2, Ifn-β, Il-6, Cxcl10, Tnf-α* and *Il-10*) in the olfactory bulb, nasal turbinates and lungs of hamsters at 4 days post-infection. The two-first principal components explained 80.5% of sample variability (n=4/group). **CD.** PCA plots. Each symbol represents one animal, colored according to the virus variant. Related to Figures 1 and 2.

**Supplementary Figure 4.**
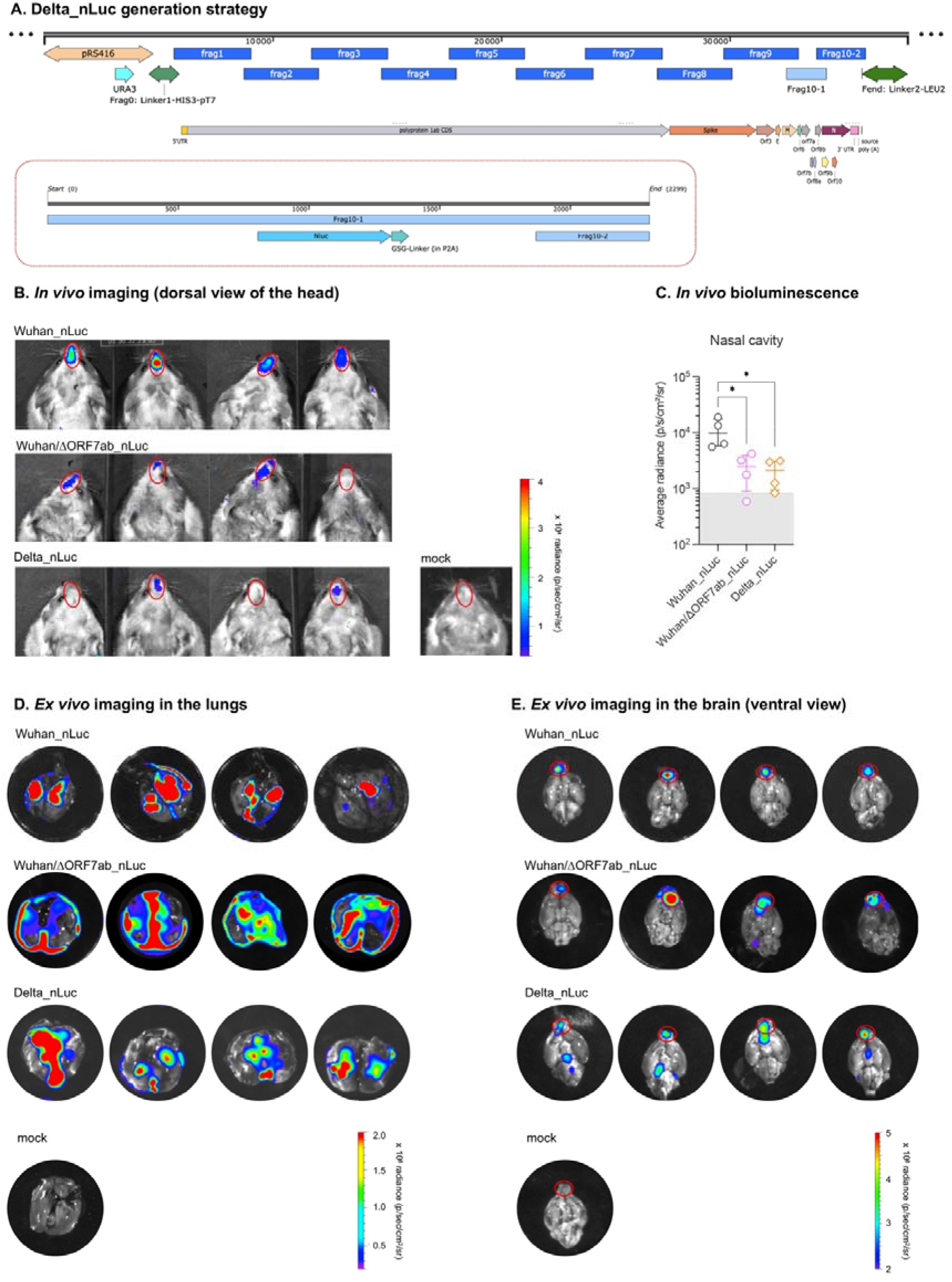
*In vivo* and *ex vivo* imaging in hamsters infected with bioluminescent recombinant SARS-CoV-2 viruses at 4 days post-infection (dpi). **A.** Delta_nLuc generation strategy. **B.** *In vivo* imaging of hamsters infected with the recombinant viruses Wuhan_nLuc and Wuhan/ΔORF7ab_nLuc (based on the original SARS-CoV-2 Wuhan) and Delta_nLuc (based on the Delta). The regions of interest (ROIs) are shown as red circles (n=4/group). **C.** *In vivo* bioluminescence values from the nasal cavity (n=4/group). Horizontal lines indicate median and the interquartile range. Kruskal-Wallis test followed by the Dunn’s multiple comparisons test (*P<0.05). The gray crosshatched area corresponds to background signals obtained from a mock-infected hamster. **DE.** *Ex vivo* imaging of lungs (D) and brains (E) from hamsters infected with Wuhan_nLuc, Wuhan/ΔORF7ab_nLuc and Delta_nLuc and mock-infected at 4 dpi. The regions of interest (ROIs) are shown as red circles in the olfactory bulbs (for the lungs and the whole brain, the values from the whole organ were acquired). Related to Figure 3.

**Supplementary Figure 5.**
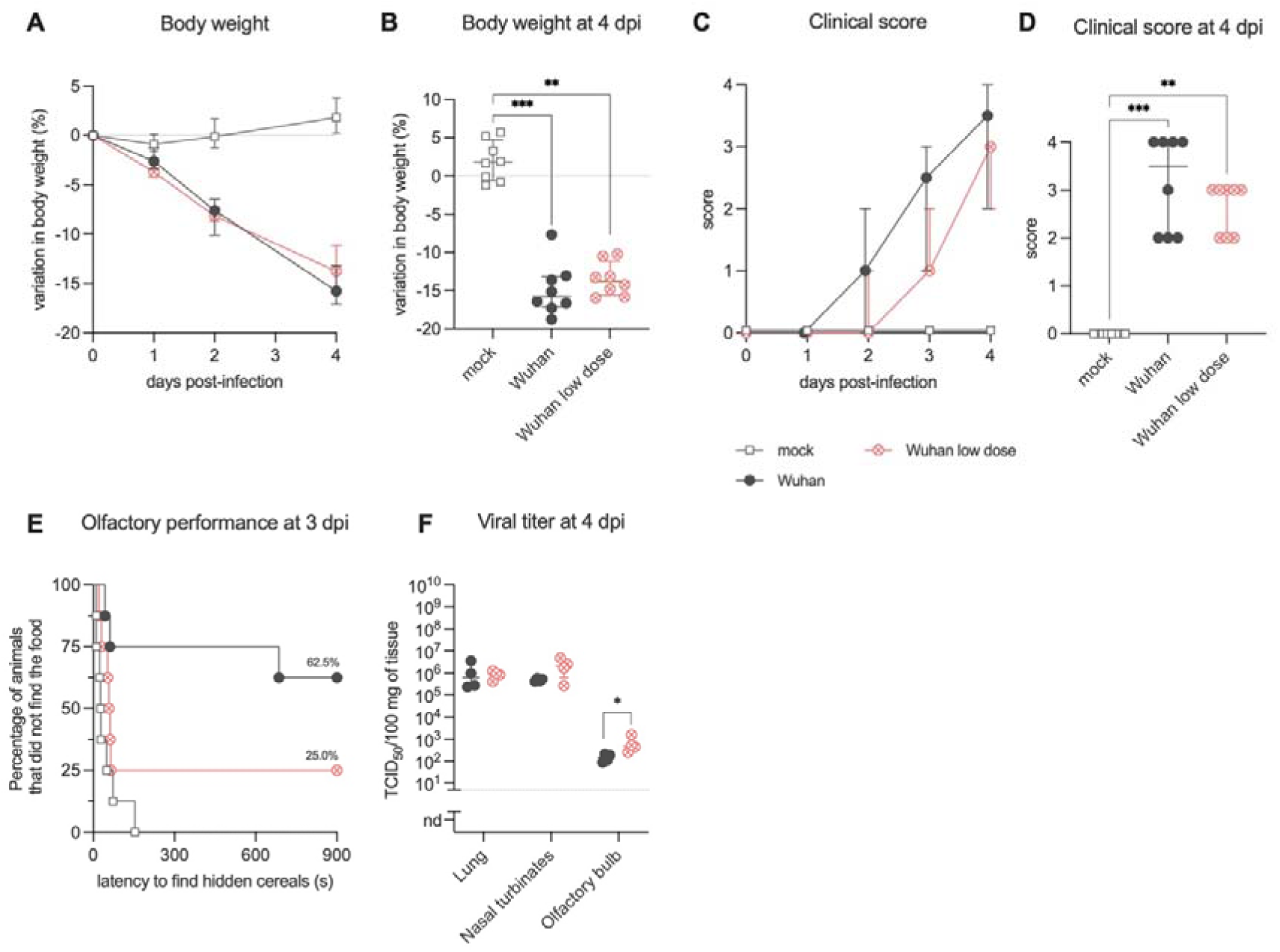
Clinical profile of hamsters inoculated with SARS-CoV-2 original virus (Wuhan) with a regular infectious dose (6x10^4^ PFU) and with a low infectious dose (6x10^2^PFU). **AB.** Body weight variation over 4 days post-infection (dpi). **CD.** Clinical score comparison in the acute phase (4 dpi). The clinical score is based on a cumulative 0–4 scale: ruffled fur; slow movements; apathy; and absence of exploration activity. Horizontal lines indicate median and the interquartile range (n=8/group). **E.** Olfactory performance measured at 3 dpi. The olfaction test is based on the hidden (buried) food finding test. Curves represent the olfactory performance of animals during the test (n=8/group). **F.** Infectious viral titers in the lung, nasal turbinates, and olfactory bulbs at 4 dpi expressed as TCID50per 100 mg of tissue. Horizontal lines indicate median and the interquartile range (n=4/group). Kruskal-Wallis test followed by the Dunn’s multiple comparisons test (*P<0.05, **P<0.01, ***P<0.001) (A-C) or Mann-Whitney test (*P<0.05) (F). Data for SARS-CoV-2 Wuhan already presented in Figures 1 and 2.

**Supplementary Figure 6.**
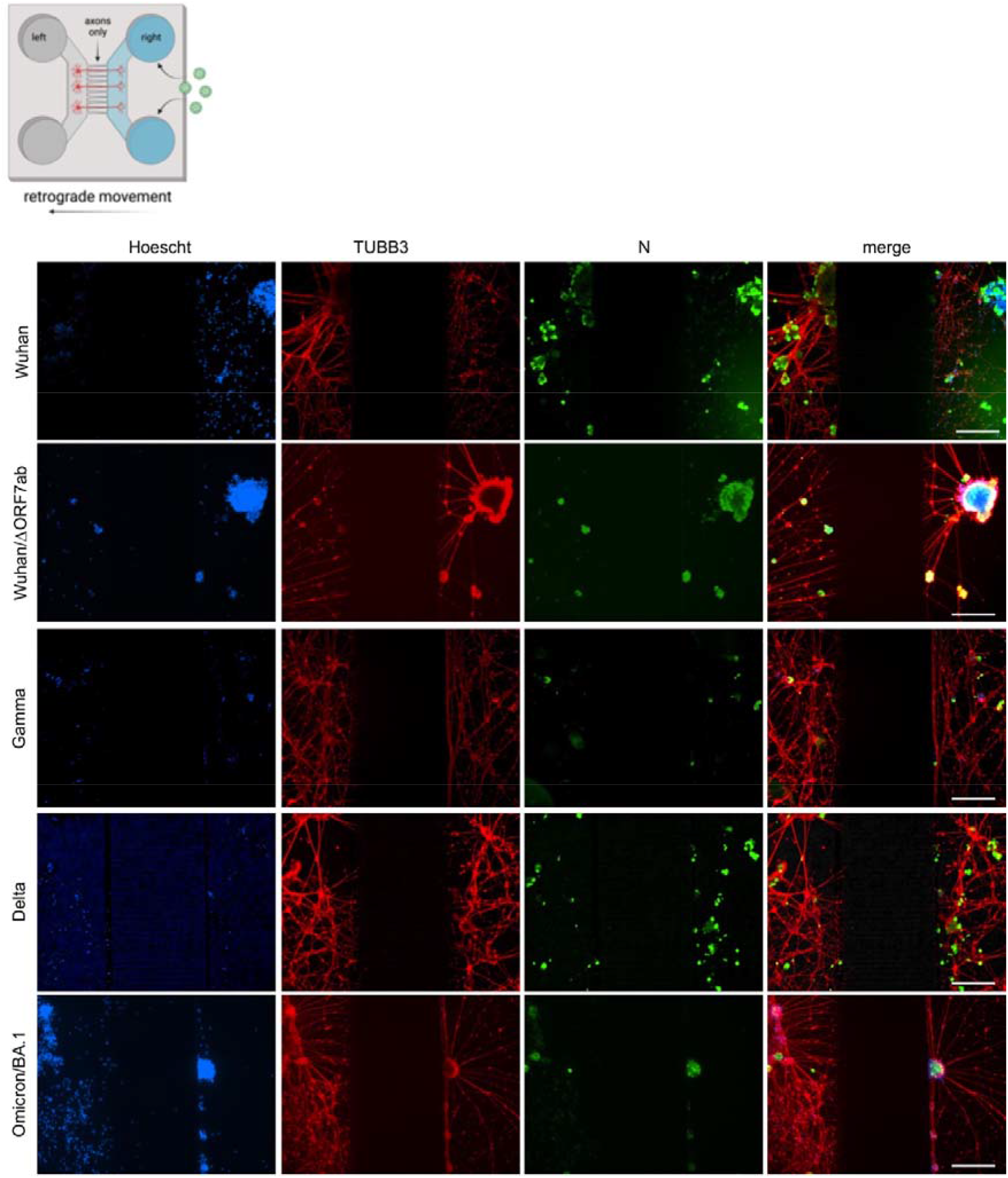
SARS-CoV-2 travels retrogradely along axons in *in vitro* neuron-epithelial networks. Schematic view of an axon diode in a microfluidic device where the left and the right chambers are connected exclusively by axons. In these images, SARS-CoV-2 was added in the right chamber: Wuhan, Wuhan/ΔORF7ab, Gamma, Delta, and Omicron/BA.1, showing infected cells in both the left and right chambers (scale bar = 200 µm). Hoechst: nuclei (blue). TUBB3: β-Tubulin III (red). N: SARS-CoV-2 nucleocapsid (green). The black zones in the central part of the photomicrographs corresponds to the microchannels, which are not accessible during immunostainings. Related to Figure 4.

**Supplementary Figure 7.**
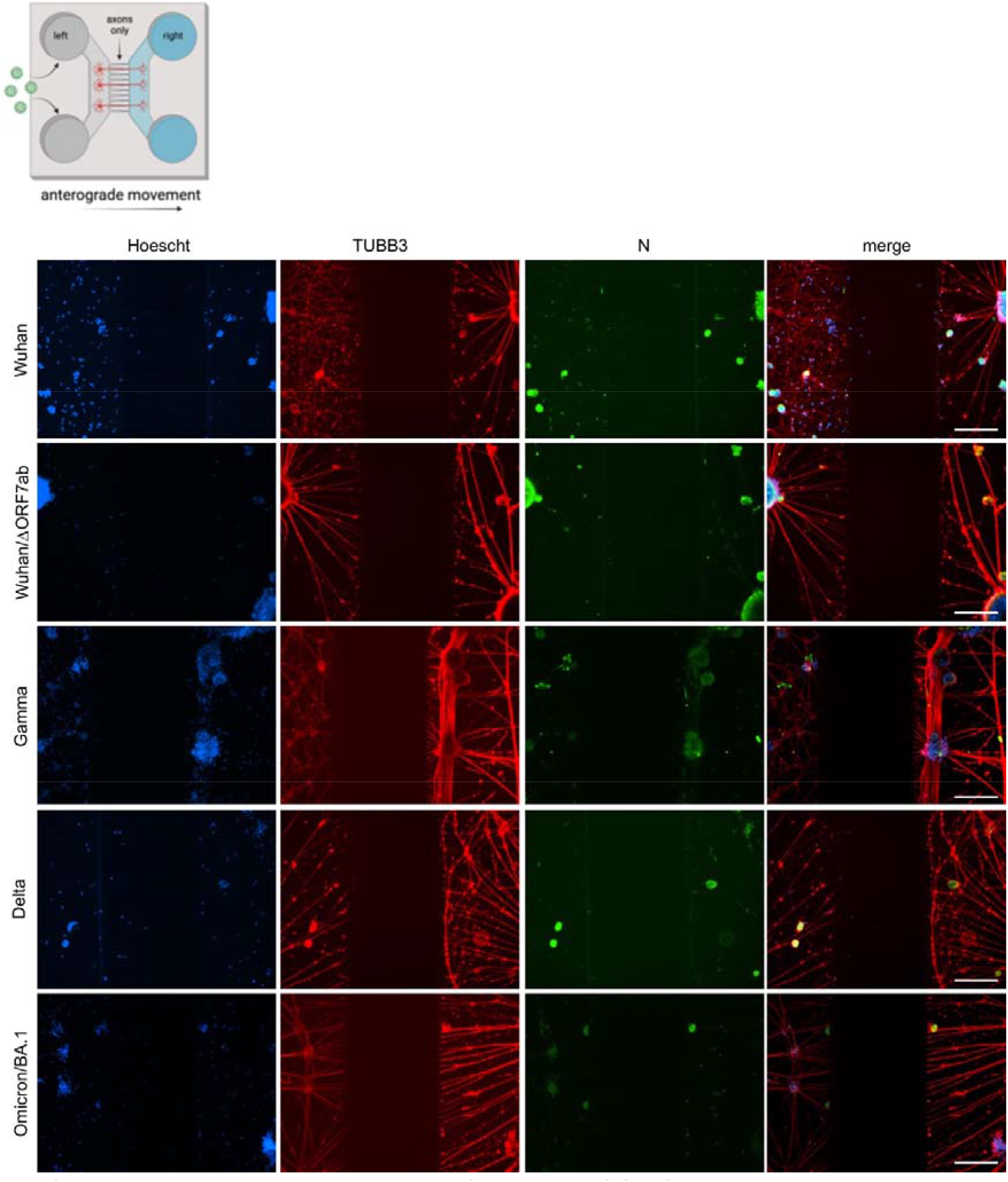
SARS-CoV-2 travels anterogradely along axons in *in vitro* neuron-epithelial networks. Schematic view of an axon diode in a microfluidic device where the left and the right chambers are connected exclusively by axons. In these images, SARS-CoV-2 was added in the left chamber: Wuhan, Wuhan/ΔORF7ab, Gamma, Delta, and Omicron/BA.1, showing infected cells in both the left and right chambers (scale bar = 200 µm). Hoechst: nuclei (blue). TUBB3: β- Tubulin III (red). N: SARS-CoV-2 nucleocapsid (green). The black zones in the central part of the photomicrographs corresponds to the microchannels, which are not accessible during immunostainings. Related to Figure 4.

**Supplementary Figure 8.**
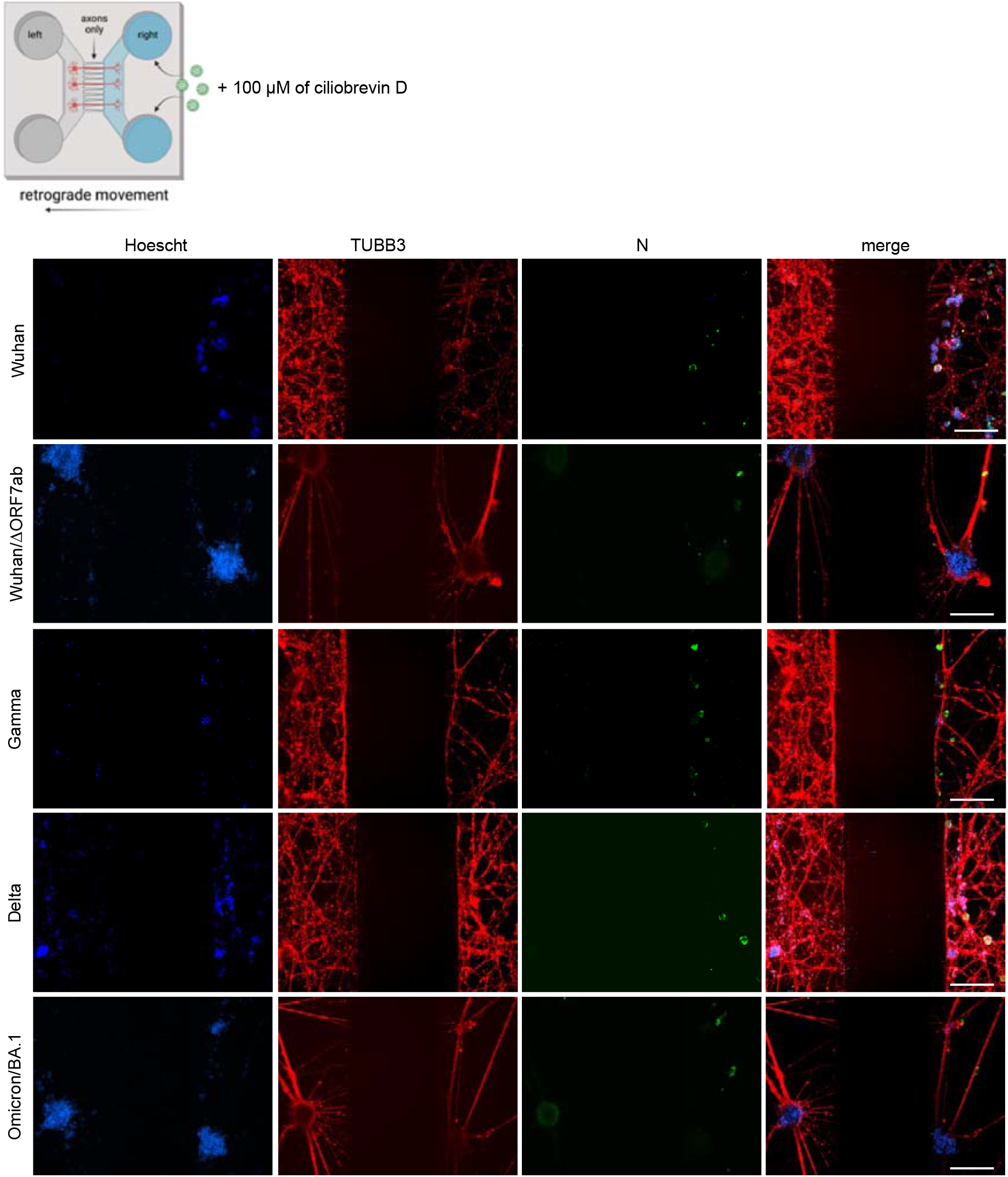
SARS-CoV-2 retrograde movement inside axons can be blocked by ciliobrevin D in *in vitro* neuron-epithelial networks. Schematic view of an axon diode in a microfluidic device where the left and the right chambers are connected exclusively by axons. In these images, 100 µM of ciliobrevin was added in both chambers and SARS-CoV-2 was added in the right chamber: Wuhan, Wuhan/ΔORF7ab, Gamma, Delta, and Omicron/BA.1, showing infected cells only in the right chambers (scale bar = 200 µm). Hoechst: nuclei (blue). TUBB3: β-Tubulin III (red). N: SARS-CoV-2 nucleocapsid (green). The black zones in the central part of the photomicrographs corresponds to the microchannels, which are not accessible during immunostainings. Related to Figure 4.

**Supplementary Figure 9.**
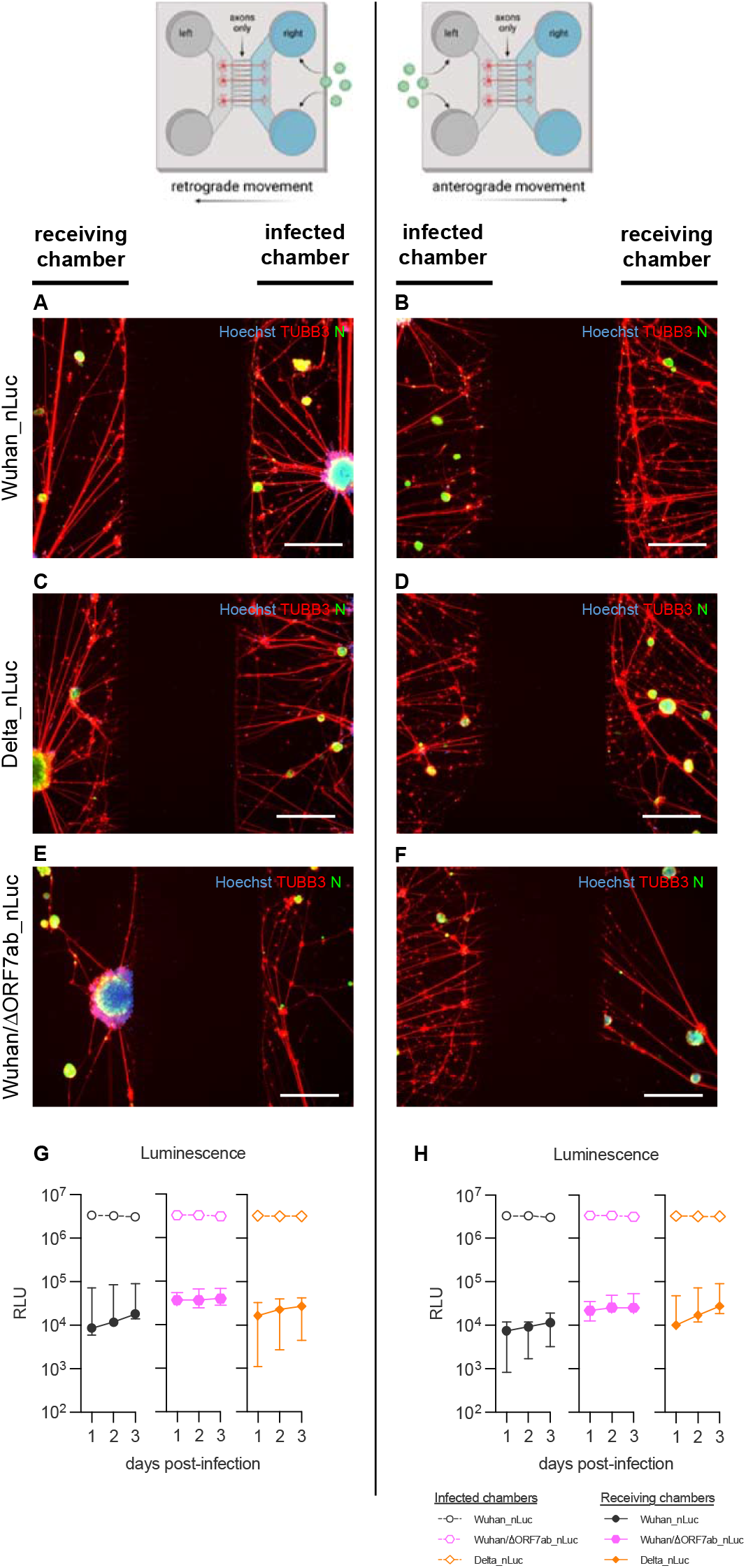
Quantification of the intra-axonal traffic of SARS-CoV-2 in *in vitro* neuron-epithelial networks. **AF.** Photomicrographs of retrograde (A,C,E) and anterograde (B, D, F) movement of the recombinant viruses Wuhan_nLuc (AB), Wuhan/ΔORF7ab_nLuc (CD) and Delta_nLuc (EF) at 3 dpi (scale bar = 200 µm). Hoechst: nuclei (blue). TUBB3: β-Tubulin III (red). N: SARS-CoV-2 nucleocapsid (green). **GH.** SARS-CoV-2 quantification by sequential luminescence measurements in the supernatants, in the retrograde (G) or anterograde (H) directions (n=3 independent replicates). Open symbols indicate the infected chambers, full-colored symbols indicate the chambers receiving the virus via the axons. RLU: relative light units. Related to Figure 4.

## Supplementary Video 1.

**Light sheet imaging in the nasal turbinates and in the olfactory bulbs.** 3D distribution of the nucleocapsid in the nostrils, including the olfactory mucosa, in the olfactory nerve and in neurons in the olfactory bulbs of a SARS-CoV-2 Wuhan infected hamster using whole head clearing. Related to Figure 3.

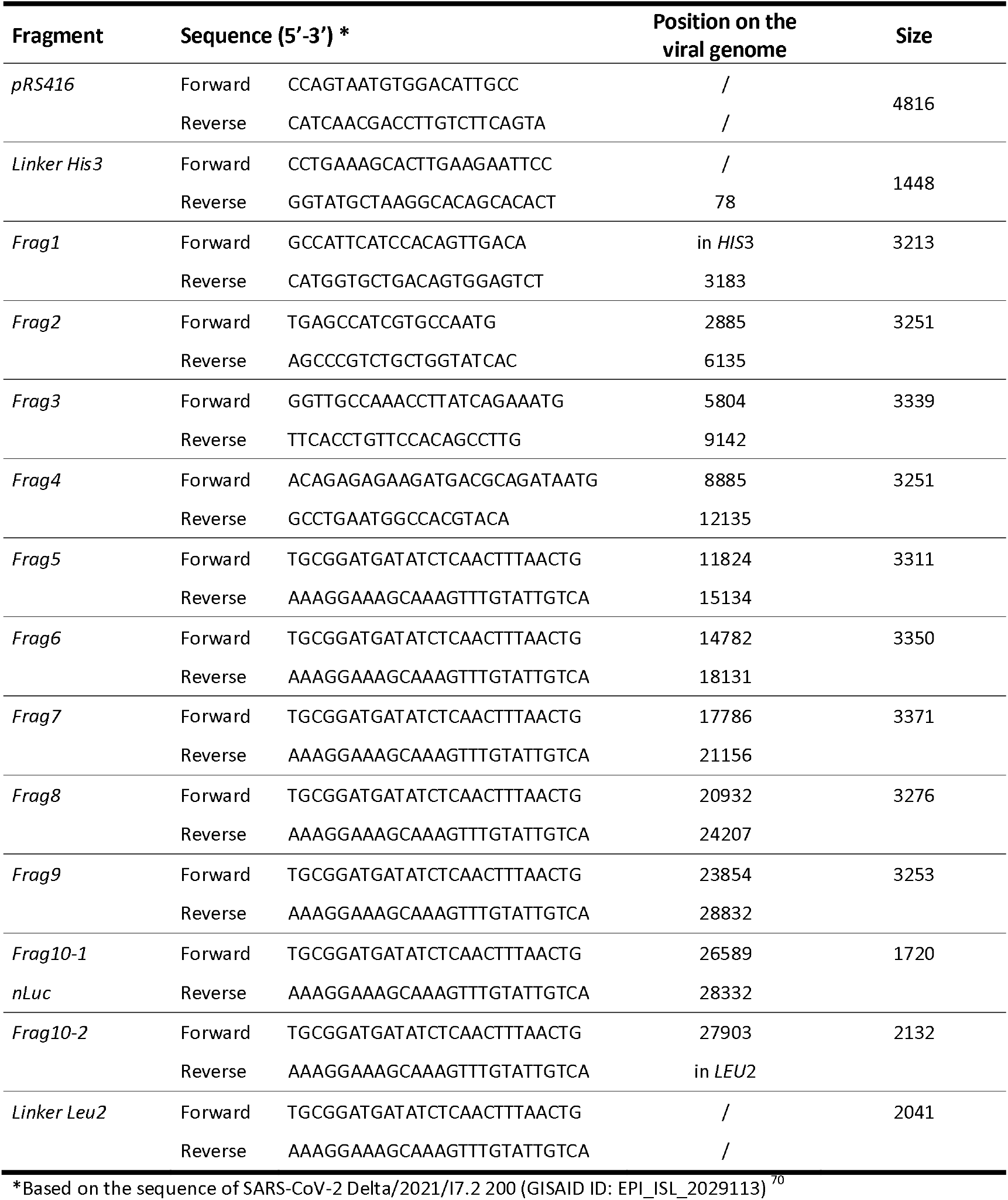
Supplementary Table 1. Primer sequences used to amplify the different fragments for yeast recombination.

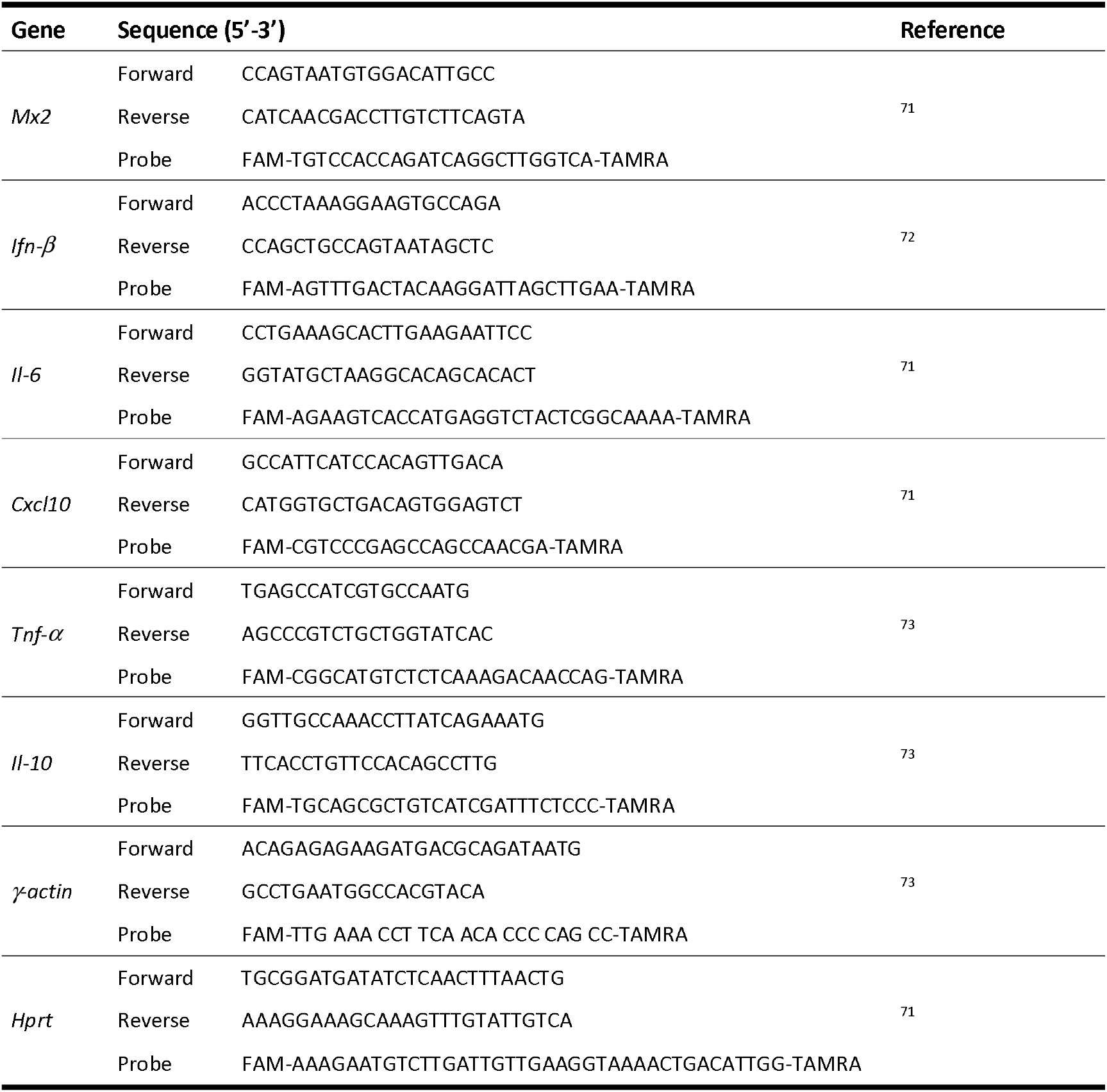
Supplementary Table 2. Primer sequences used for qPCR in the golden hamster tissues.

### Materials and methods

#### Ethics

All animal experiments were performed according to the French legislation and in compliance with the European Communities Council Directives (2010/63/UE, French Law 2013– 118, February 6, 2013) and according to the regulations of Institut Pasteur Animal Care Committees. The Animal Experimentation Ethics Committee (CETEA 89) of the Institut Pasteur approved this study (200023; APAFIS#25326-2020050617114340 v2) before experiments were initiated. Hamsters were housed by groups of 4 animals in isolators and manipulated in class III safety cabinets in the Pasteur Institute animal facilities accredited by the French Ministry of Agriculture for performing experiments on live rodents. All animals were handled in strict accordance with good animal practice.

#### SARS-CoV-2 virus and variants

The isolate BetaCoV/France/IDF00372/2020 (EVAg collection, Ref SKU: 014V-03890) was kindly provided by the National Reference Centre for Respiratory Viruses hosted by Institut Pasteur (Paris, France) and headed by Pr. Sylvie van der Werf. The human sample from which strain hCoV-19/Japan/TY7-501/2021 (Brazilian variant, JPN [P.1]) was supplied by the Japanese National Institute of Infectious Diseases (Tokyo, Japan). The isolate SARS-CoV-2 Delta/2021/I7.2 200 (Indian variant, GISAID ID: EPI_ISL_2029113) and the isolate SARS-CoV-2 Omicron/B.1.1.529 (Omicron BA.1 variant, GISAID ID: EPI_ISL_6794907) were kindly provided by the Virus and Immunity Unit hosted by Institut Pasteur and headed by Olivier Schwartz ^70, 74^. Viral stocks were produced on Vero-E6 cells infected at a multiplicity of infection of 1x10^-4^ PFU/cell. The virus was harvested 3 days post-infection, clarified and then aliquoted before storage at -80°C. Viral stocks were titrated on Vero-E6 cells by classical plaque assays using semisolid overlays (Avicel, RC581-NFDR080I, DuPont) ^75^.

Recombinant viruses expressing Nanoluciferase (nLuc) as a reporter gene were obtained by reverse genetics. Wuhan_nLuc was derived from the clone synSARS-CoV-2-GFP-P2A-ORF7a (clone #41) generated as described ^76–80^, where the GFP reporter gene was replaced by the nLuc one. The cloning strategy was optimized to build a complete cDNA of the Delta_nLuc virus on the same backbone than the isolate EPI_ISL_2029113. SARS-CoV-2 Wuhan/ΔORF7ab was obtained using the same reverse genetic system by replacing the complete ORF7ab sequence by that of GFP or by nLuc (Wuhan/ΔORF7ab_nLuc).

#### Generation of the complete genome of SARS- CoV-2/Delta_nLuc virus

The viral genome was divided in 11 overlapping fragments around 3kb each. Those fragments and two fragments coding for different yeast-specific selection genes (His3 and Leu2) were recombined in the yeast centromere plasmid pRS416 to generate the complete genome clone of recombinant virus. The T7 promotor was placed just before the 5’UTR sequence and a unique restriction site EAG1 was placed just after the poly(A) tail.

Viral fragments were obtained either by RT-PCR on RNA extracted from Vero-E6 cells infected by Delta using the SuperScript IV VILO Master Mix (11756050, Thermofisher) according to the manufacturer protocol, or using synthetic genes (GeneArt, Life Technologies). All PCR amplifications were performed using Phusion™ High-Fidelity DNA Polymerase (F530, Thermofisher). The PCR product were cloned in Topo-TA vector and their sequences controlled by sanger sequencing. The nLuc gene was inserted upstream ORF7a and separated from it using a GSG linker and a P2A peptide sequence to mediate cleavage between the reporter and the viral protein. It was directly synthetized in the F10-1 fragment.

#### Yeast recombination

Recombination of the 4 yeast specific fragments and the 11 viral fragments was performed on *S. cerevisiae* BY4741. A yeast culture was carried out from an overnight pre culture in YPDA medium (2 mL of pre-culture in 50 mL of YPDA medium heated to 30° C) until the exponential growth phase was reached, with an optical density corresponding to approximately 10^8^ cells/mL. The yeasts were harvested by centrifugation of 50 mL of culture (5,000 rpm at 4°C during 5 min) before being washed a first time in sterile solution of Tris-EDTA and lithium Acetate TE/LiAc (Tris-HCl pH 7.5 10 mM; EDTA pH 7.5 0.1 mM; LiAc 100 mM). Cells were resuspended in 500 μL of TE/LiAc solution (2.10^9^ cells/mL). The yeast suspension was then incubated at 30° C without stirring during 30 min. During this time, a mix of all DNA fragments in an equimolar proportion was prepared. For each recombination condition, 50 μL of competent cells (10^8^cells) were resuspended in a final volume of 300 μL TE/LiAc/PEG solution (100 mM Tris-HCl pH 8; 100 mM Lithium Acetate; 40% v/v PEG 3350) with the DNA mix to be recombined and 50 μg of denaturated salmon sperm DNA (31149, Sigma) and then cooled on ice for 5 min. The mixture was incubated at 30°C during 30 min without agitation before undergoing a heat shock at 42°C during 20 min. After cooling on ice, the yeasts were pelleted by centrifugation at 2,000 rpm at RT during 5 min and then resuspended and incubated with 500 μL of 5 mM CaCl2at RT for 10 min. CaCl2was washed by centrifugation at 2,000 rpm at RT during 5 min and cells were resuspended in 200 μL of sterile water before being plated on selective synthetic minimal medium lacking histidine, Uracyl and Leucine (SD-His-Ura-Leu-), and incubated at 30°C for 3 days. For each transformation, ten different clones were checked by multiplex PCR (206143, Qiagen) on rapid DNA preparations carried out by the Chellex technique as already described ^76^ using specific primers (Table S1) to control the presence of the different fragments in the expected orientation. The yeast colony of interest was sub-cultured in 200 mL of SD-His-Ura-Leu-medium and the plasmid extracted using the Qiagen midi prep kit (12143, Qiagen) following manufacturer’s instructions.

#### Rescue of the virus

The plasmid was linearized by EagI-HF (R3505, NEB) digestion, purified by a phenol chloroform process and transcribed *in vitro* into RNA using the T7 RiboMaxTM Large Scale RNA Production System (P1300, Promega) and the m7G(5’)ppp(5’)G RNA Cap Structure Analog (#S1411, NEB) kits. Approximately 5 μg of linear DNA were used as template in a reaction volume of 50 μL comprising: 10 μL of 5X T7 transcription buffer; 5 μL of m7G(50)ppp(50)G RNA cap analog structure at 30 mM; 0.75 μL of 100 mM GTP; 3.75 μL of each nucleotide type ATP, CTP; 100 mM UTP; and 5 μL of RNasin T7 RNA polymerase enzyme. The reaction was incubated at 30°C during 3 h then the template DNA was digested at 37°C during 20 min by adding 2 μL of the enzyme RQ1 RNase free DNase. The synthesized RNA was finally purified using a classical phenol chloroform method and precipitated. Twelve μg of complete viral mRNA and 4 μg of plasmid encoding the viral nucleoprotein (N) gene were electroporated on 8x10^6^ Vero-E6 cells resuspended in 0.8 mL of Mirus Bio™ Ingenio™ Electroporation Solution (MIR50114) using the Gene Pulser Xcell Electroporation System (1652660, Biorad) with a pulse of 270V and 950 μF. The cells were then transferred to a T75 culture flask with 15 mL of DMEM supplemented with 2% FCS (v/v). The electroporated cells were incubated at 37°C, 5% CO2for several days until the cytopathic effect (CPE) was observed. Then, the supernatant was harvested, aliquoted and frozen at -80°C until titration. The viral sequence was controlled by NGS.

#### Viral growth curves

Vero-E6 cells (ATCC CRL-1586) were used to assess the replication kinetics of different SARS-CoV-2 variants and recombinant viruses. The cells were maintained in Dulbecco’s modified culture medium (DMEM, 31966-021, Gibco) supplemented with 5% fetal calf serum, at 37bC in 5% CO2. On the day before infection, 1x10^6^VeroE6 cells/well were plated onto 6-well plates. On day of infection, the cells were infected with SARS-CoV-2 virus at a multiplicity of infection (MOI) of 0.01, in independent triplicates, and incubated during 24-, 48- or 72-hours post infection. The supernatants were collected at each time-point and frozen at -80°C. Viral titers were obtained by classical TCID50 method on Vero-E6 cells after 72 hours post-infection ^81^.

#### SARS-CoV-2 model in golden Syrian hamsters

Male golden Syrian hamsters (*Mesocricetus auratus;* RjHan:AURA) of 5-6 weeks of age (average weight 60-80 grams) were purchased from Janvier Laboratories and handled under specific pathogen-free conditions. The animals were housed and manipulated in isolators in a biosafety level-3 facility, with *ad libitum* access to water and food. Before any manipulation, animals underwent an acclimation period of one week. Animals were anesthetized with an intraperitoneal injection of 200 mg/kg ketamine (Imalgène 1000, Merial) and 10 mg/kg xylazine (Rompun, Bayer), and 100 µL of physiological solution containing 6x10^4^ PFU of SARS-CoV-2 (wild-type or recombinant) was administered intranasally to each animal (50 µL/nostril). Mock-infected animals received the physiological solution only. Infected and mock infected hamsters were housed in separate isolators and were followed-up daily during four days at which the body weight and the clinical score were noted. The clinical score was based on a cumulative 0-4 scale: ruffled fur, slow movements, apathy, absence of exploration activity. At day 3 post-infection (dpi), animals underwent a food finding test to assess olfaction as previously described ^14, 82^. Briefly, 24 hours before testing, hamsters were fasted and then individually placed into a fresh cage (37 x 29 x 18 cm) with clean standard bedding for 10 minutes. Subsequently, hamsters were placed in another similar cage for 2 minutes when about 5 pieces of cereals were hidden in 1.5 cm bedding in a corner of the test cage. The tested hamsters were then placed in the opposite corner and the latency to find the food (defined as the time to locate cereals and start digging) was recorded using a chronometer. The test was carried out during a 15 min period. As soon as food was uncovered, hamsters were removed from the cage. One minute later, hamsters performed the same test but with visible chocolate cereals, positioned upon the bedding. The tests were realized in isolators in a Biosafety level-3 facility that were specially equipped for that. At 4 dpi, animals were euthanized with an excess of anesthetics (ketamine and xylazine) and exsanguination ^83^, and samples of nasal turbinates, lungs and olfactory bulbs were collected and immediately frozen at -80°C. Fragments of lungs were also collected and fixed in 10% neutral buffered formalin.

#### SARS-CoV-2 detection in golden hamsters’ tissues

Frozen lung fragments, nasal turbinates and olfactory bulbs were weighted and homogenized with 1 mL of ice-cold DMEM supplemented with 1% penicillin/streptomycin (15140148, Thermo Fisher) in Lysing Matrix M 2 mL tubes (116923050- CF, MP Biomedicals) using the FastPrep-24™ system (MP Biomedicals), and the following scheme: homogenization at 4.0 m/s during 20 sec, incubation at 4°C during 2 min, and new homogenization at 4.0 m/s during 20 sec. The tubes were centrifuged at 10.000 x g during 2 min at 4°C, and the supernatants collected. Viral titers were obtained by classical TCID50 method on Vero-E6 cells after 72 hours post-infection ^81^. Viral RNA loads were obtained by the quantification of genomic and sub genomic SARS-CoV-2 RNA based on the E gene ^84^. Briefly, 125 µL of the supernatants were homogenized with 375 µL of Trizol LS (10296028, Invitrogen) and total RNA was extracted using the Direct-zol RNA MicroPrep Kit (R2062, Zymo Research: nasal turbinates and olfactory bulbs) or MiniPrep Kit (R2052, Zymo Research: lung). We used the Taqman one-step qRT-PCR (Invitrogen 11732-020) in a final volume of 12.5bμL per reaction in 384-wells PCR plates using a thermocycler (QuantStudio 6 Flex, Applied Biosystems). Briefly, 2.5bμL of RNA were added to 10bμL of a master mix containing 6.25bμL of 2X reaction mix, 0.2 µL of MgSO4(50 mM), 0.5 µL of Superscript III RT/Platinum Taq Mix (2 UI/µL) and 3.05bμL of nuclease-free water containing 400 nM of primers and 200 nM of probe. To detect the genomic RNA, we used the E_sarbeco primers and probe (E_Sarbeco_F1 5b-ACAGGTACGTTAATAGTTAATAGCGT-3’; E_Sarbeco_R2 5b- ATATTGCAGCAGTACGCACACA-3’; E_Sarbeco_Probe FAM-5b-ACACTAGCCATCCTTACTGCGCTTCG-3’-TAMRA). The detection of sub-genomic SARS-CoV-2 RNA was achieved by replacing the E_Sarbeco_F1 primer by the CoV2sgLead primer (CoV2sgLead-Fw 5b- CGATCTCTTGTAGATCTGTTCTC-3’). A synthetic gene encoding the PCR target sequences was ordered from Thermo Fisher Scientific. A PCR product was amplified using Phusion™ High-Fidelity DNA Polymerase (Thermo Fisher Scientific) and *in vitro* transcribed by means of Ribomax T7 kit (Promega). RNA was quantified using Qubit RNA HS Assay kit (Thermo Fisher scientific), normalized, and used as a standard to quantify RNA absolute copy number. The amplification conditions were as follows: 55°C for 20bmin, 95°C for 3 minutes, 50 cycles of 95°C for 15bs and 58°C for 30 s; followed by 40°C for 30 s.

#### Transcriptomics analysis in golden hamsters’ tissues

RNA preparations from lungs, nasal turbinates and olfactory bulbs collected at 4 dpi were submitted to RT-qPCR. Briefly, RNA was reverse transcribed to first strand cDNA using the SuperScript™ IV VILO™ Master Mix (11766050, Invitrogen). qPCR was performed in a final volume of 10bμL per reaction in 384-well PCR plates using a thermocycler (QuantStudio 6 Flex, Applied Biosystems). Briefly, 2.5bμL of cDNA (12.5 ng) were added to 5bμL of Taqman Fast Advanced master mix (444457, Applied Biosystems) and 2.5bμL of nuclease-free water containing golden hamster’s primer pairs (Table S2). The amplification conditions were as follows: 95°C for 20 s, and 45 cycles of 95°C for 1bs and 60°C for 20 s. The *γ-actin* and the *Hprt* (hypoxanthine phosphoribosyl-transferase) genes were used as reference. Variations in gene expression were calculated as the n-fold change in expression in the tissues from the infected hamsters compared with the tissues of the mock-infected group using the 2^-^*^ΔΔ^*^Ct^method ^85^.

#### Sample staining and iDISCO+ clearing

For iDISCO+, three male and three female hamsters were infected with 6x10^4^PFU of SARS-CoV-2 Wuhan and followed up as described above. At 4 dpi, animals were anesthetized with an intraperitoneal injection of ketamine (200 mg/kg; Imalgène 1000, Merial) and xylazine (10 mg/kg; Rompun, Bayer), and we performed a transcardial perfusion with DPBS containing heparin (5×10^3^U/mL) followed by 4% neutral-buffered formaldehyde. The whole heads were collected and stored in 4% neutral-buffered formaldehyde during one week before analysis. SARS-CoV-2 nucleocapsid was detected *in toto* in the snout and brain of infected hamsters using the iDISCO+ protocol previously published ^86^ with minimal modifications. All buffers were supplemented with 0,01% of sodium azide (Sigma-Aldrich). The snout, containing the olfactory mucosa and the olfactory nerve, and the brain were dissected and processed separately to optimize tissue manipulation.

All samples were first dehydrated in methanol (Sigma-Aldrich) using 20-40-60-80-100-100% dilutions in distilled water (1h to 2h each concentration). Complex lipid removal was achieved incubating the samples in a 2:1 mixture of dichloromethane (Sigma-Aldrich) and methanol overnight. Samples were then washed twice in methanol 100% and bleached overnight using a H2O25% solution in methanol. Then all samples were rehydrated using a methanol series (80-60- 40-20%). To increase bone permeability, the snouts were decalcified by incubation in the Morse’s solution (formic acid 45% and sodium citrate 20%). Finally, all the samples (brains and snouts) were washed in PBS, then PBS-T (PBS with 0,2% Triton X-100 [Sigma-Aldrich]) and incubated in permeabilization buffer (20% DMSO [Sigma-Aldrich] and 2.3% glycine [Sigma-Aldrich] in PBS-T) at 37°C overnight. Before immunostaining, samples were blocked (0,2% Gelatin [Sigma-Aldrich] in PBS-T) at 37°C for 24h. To reveal SARS-CoV-2 and vasculature, three primary antibodies were combined: Rabbit anti-SARS CoV2 nucleocapsid antibody (GTX135357, GeneTex) diluted 1:1000; Goat anti-CD31 (AF3628, R&D Systems) diluted 1:300; and Rat anti-Podocalyxin (MAB1556, R&D Systems) diluted 1:1000. After a 2-weeks incubation in primary antibody, the samples were washed (PBS supplemented with 0,2% tween-20 [Sigma-Aldrich] and 2U/mL heparin [Sigma-Aldrich]) and incubated 10 days with the following secondary antibodies: Donkey anti-Rabbit Alexa 555 (A-31572, Thermo Fisher Scientific), Donkey anti-Goat Alexa 647 (A-21447, Thermo Fisher Scientific) and Chicken anti-Rat Alexa 647 (A-21472, Thermo Fisher Scientific) all diluted at 1:500. All antibodies were diluted in blocking solution and incubated at 37°C with gentle shacking. After immunostaining, the samples were washed, dehydrated in methanol (20-40-60-80-100-100%), incubated for 3h in a 2:1 mixture of dichloromethane and methanol, washed twice (15 min each) in dichloromethane 100% and cleared by immersion in dibenzyl ether.

#### Light sheet imaging

Brain and snout samples were imaged using a LaVision Ultramicroscope II equipped with infinity-corrected objectives, laser lines OBIS-561nm 100mW and OBIS-639nm 70mW, and 595/40 and 680/30 filters for Alexa 555 and Alexa 647 respectively. The olfactory bulbs were imaged with a 4X 0,35NA objective, using a laser NA of 0,3 and a step size of 2µm obtaining images with 1,63x1,63x2µm/pixel resolution. The snouts were imaged with a 1,3X objective, adjusting the laser NA to 0,3 and step size of 5µm obtaining images with a 5x5x5µm/pixel resolution. All acquisitions were done with LaVision BioTec ImSpector Software. The 3D stacks obtained were analyzed using Bitplane Imaris 9.2 (Oxford instruments).

#### *In vivo* and *ex vivo* bioluminescence imaging

At 4 dpi, animals infected with SARS-CoV- 2/Wuhan_nLuc, Wuhan/ΔORF7ab_nLuc and SARS-CoV-2/Delta_nLuc were anesthetized with an intraperitoneal injection of ketamine (200 mg/kg; Imalgène 1000, Merial) and xylazine (10 mg/kg; Rompun, Bayer). Next, 0.45 µmoles of the nLuc substract (Nano-Glo® *in vivo* substrate, CS320501, Promega) fluorofurimazine (FFz) were injected interperitoneally and the animals were placed in a confinement box and imaged using an IVIS® Spectrum *In Vivo* Imaging System (PerkinElmer) within 5 minutes of FFz injection. Two-dimensional bioluminescence images were recorded, and photon emission was quantified (p/s/cm^2^/sr) in a region of interest defined using Living Image software (PerkinElmer). After *in vivo* imaging, animals were euthanized, the lungs and the brains were collected and quickly placed in a six-wells plate. The plates were placed in a confinement box and imaged as described above.

#### Histopathology and Immunohistochemistry

Lung fragments fixed 7 days in 10% neutral-buffered formalin were embedded in paraffin. Four-µm-thick sections were cut and stained with hematoxylin and eosin staining. IHC were performed on Leica Bond RX using anti SARS Nucleocapsid Protein antibody (NB100-56576, Novus Biologicals) and biotinylated goat anti-rabbit Ig secondary antibody (E0432, Dako, Agilent). Slides were then scanned using Axioscan Z1 slide scanner (Zeiss) and images were analyzed with the Zen 2.6 software (Zeiss).

#### Neuron-epithelial networks in microfluidic chambers

Human neural stem cells (hNSC, ENStem-A, SCC003, EMD-Millipore) were maintained in geltrex-coated 6-well plates (A1413302, Gibco) in a density of 4.5x10^5^ cells/cm^2^in CTS Knock-out DMEM/F12 medium (A1370801, Gibco) supplemented with 2% StemPro neural supplement (A1050801, Gibco), 2 mM glutamax (35050038, Gibco), 20 ng/mL FGFb (PHG0026, Gibco) and 20 ng/mL EGF (PHG0311, Gibco) at 37°C in 5% CO2. A549-ACE2-TMPRSS2 cells (kindly provided by Pr. Olivier Schwartz, Institut Pasteur) were maintained in F12K Nut Mix medium (21127022, Gibco) supplemented with 10% bovine calf serum and 10 µg/mL blasticidine (12172530, Gibco) at 37°C in 5% CO2. To generate the neuron-epithelial networks, we used microfluidic chips purchased from MicroBrain Biotech (Brainies™, Cat#: MBBT5; Marly le Roi, France). Brainies™ MBBT5 is a chip containing 4 neuronal diodes. One neuronal diode includes 2 rectangular culture chambers (volume ∼1 µL) each connected to 2 reservoirs and separated by a series of 500 µm-long asymmetrical micro-channels (3 µm high, tapering from 15 µm to 3 µm) ^33^. Brainies™ were coated with poly-L-ornithine (P4957, Sigma-Aldrich) and laminin (L2020, Sigma-Aldrich) and 1x10^5^hNSCs were seeded in both the right and the left chambers and incubated at 37°C in 5% CO2for 24 hours. The medium was then replaced by a neuronal differentiation medium composed of neurobasal medium (10888022, Gibco) supplemented with 2% B27 supplement (17504044, Gibco), 1% CultureOne supplement (A3320201, Gibco), 2 mM glutamax (35050038, Gibco), and 200 µM ascorbic acid (A4403, Sigma Aldrich) and incubated at 37°C in 5% CO_2_ for 14 days, replacing half of the medium every other day. Finally, 2x10^4^A549-ACE2- TMPRSS2 cells were added over the neurons in both the right and the left chambers in neuronal differentiation medium at 37°C in 5% CO2for 48 hours. To assess the SARS-CoV-2 retrograde axonal movement, the right chambers were infected at a MOI of 1 of SARS-CoV-2 Wuhan, Wuhan/ΔORF7ab, Gamma, Delta, or Omicron/BA.1, and incubated at 37°C in 5% CO2for 72 hours. To assess the SARS-CoV-2 anterograde axonal movement, the left chambers were infected in the same abovementioned conditions. To prevent passive diffusion of assemblies, hydrostatic pressure was applied by adding medium excess in the reservoirs of recipient chambers ^87^. To block the axonal retrograde transport, 100 µM of ciliobrevin D (250401, Calbiochem) was added in the medium during infection and throughout the incubation period. The cells were then fixed with 4% PFA (15444459, Thermo Scientific), washed in PBS, permeabilized with 0.5% Triton X-100 during 10 minutes, washed once in PBS, blocked with 10% normal goat serum (10000C, Invitrogen) during 30 minutes followed by overnight incubation at 4°C with primary antibodies: mouse anti-β-Tubulin III (neuronal) antibody (1:1000, T8578, Sigma-Aldrich) and rabbit anti-SARS-CoV-2 nucleocapsid antibody (1:1000, GTX135361, GeneTex). The cells were then washed in PBS and incubated with the following secondary antibodies: goat anti-mouse AlexaFluor 647 (1:1000, A21235, Invitrogen) and goat anti-rabbit AlexaFluor 546 (1:1000, A11035, Invitrogen) for 2 hours at 4°C. The cells were then washed, the nuclei were stained with 20 µM Hoechst 33342 (62249, Thermo Scientific) and stored in PBS. Images were obtained using the EVOS FL imaging system (Invitrogen) with the objectives 10x or 20x and the fluorescence cubes DAPI, RFP and CY5.5. Using the same described method, neuron-epithelial networks in microfluidics devices were also infected with the recombinant viruses Wuhan_nLuc, Wuhan/**Δ**ORF7ab_nLuc, and Delta_nLuc at a MOI of 0.5 (maximum volume limitation due to stock viral titer). The nanoluciferase activity in the supernatants was evaluated sequentially at 1, 2 and 3 dpi. Briefly, 20 µL of supernatant was mixed with 20 µL of nLuc substrate (N1110, Promega) in a white 96- wells plate and the luminescence, expressed as Relative light Unit (RLU), was acquired during 100 ms using the VICTOR Nivo Plate reader (Perkin Elmer).

#### Statistics

Statistical analysis was performed using Prism software (GraphPad, version 9.0.0, San Diego, USA), with *p* < 0.05 considered significant. Quantitative data was compared across groups using Log-rank test, two-tailed Mann-Whitney test or Kruskal-Wallis test followed by the Dunn’s multiple comparisons test. Multivariate statistical analyses on clinical parameters and on tissue inflammation were achieved using Principal Component Analysis. Randomization and blinding were not possible due to pre-defined housing conditions (separated isolators between infected and non-infected animals). *Ex vivo* analyses were blinded (coded samples).

## Acknowledgements

The SARS-CoV-2 strain was supplied by the National Reference Centre for Respiratory Viruses hosted by Institut Pasteur (Paris, France) and headed by Pr. Sylvie van der Werf. The human sample from which strain 2019-nCoV/IDF0372/2020 was isolated has been provided by Pr. X. Lescure and Pr. Y. Yazdanpanah from the Bichat Hospital (Paris, France). The human sample from which strain hCoV-19/Japan/TY7-501/2021 (Gamma variant, JPN [P.1]) was supplied by the Japanese National Institute of Infectious Diseases (Tokyo, Japan). The isolate SARS-CoV-2 Delta/2021/I7.2 200 (Delta variant, GISAID ID: EPI_ISL_2029113), the isolate SARS-CoV-2 Omicron/B.1.1.529 (Omicron BA.1 variant, GISAID ID: EPI_ISL_6794907) and the A549-ACE2-TMPRSS2 cells were supplied by the Virus and Immunity Unit hosted by Institut Pasteur and headed by Pr. Olivier Schwartz. We thank Sanjay Vashee from the J. Craig Venter Institute (Rockville, MD, USA) for the YCpBAC-his3 plasmid. This work was supported by Institut Pasteur’s Task Force SARS-CoV-2 (NeuroCovid Project), by SARS-CoV-2 joint call Institut Pasteur - Paris Brain Institute (CoVessel Project), by Institut Pasteur’s Programme Fédérateur de Recherche 1 (PFR-1 - Reverse Genetics). G.D.M. acknowledges funding from the Fondation pour la Recherche Médicale (grant ANRS MIE202112015304). V.P. is recipient of a fellowship from the European Union’s Horizon 2020 Framework Programme for Research and Innovation under Specific Grant Agreement No. 945539 (Human Brain Project SGA3). F.A. is recipient of a fellowship from Institut Pasteur’s Programme Fédérateur de Recherche 5 (PFR-5 - Functional Genomics of the Viral Cycle). A.C. acknowledges funding from the Institut Pasteur’s 2022-2023 Brain Axis SRA3 M2 Master Student Call. E.S.L and R.K acknowledge support from the Institut Pasteur’s Task Force (project SABSOS). E.S.L acknowledges funding from the INCEPTION programme (Investissements d’Avenir grant ANR-16-CONV-0005). We would like to acknowledge Yves Jacob for the fruitful discussions about SARS-CoV-2 ORF7. We thank Etienne Jacotot for his insights on neuronal cultures in microfluidic chambers, as well as Bernadette Bung from MicroBrain Biotech. We also thank Johan Bedel for the help with histopathology, and Emeline Perthame for her insights on statistical analyses. Part of this work was performed at the UtechS Photonic BioImaging (PBI) platform supported by Institut Pasteur and by Région Ile-de-France (program DIM1Health).

## Author Contributions

Conceptualization: G.D.M., F.L., and H.B.

Methodology: G.D.M., R.K., V.T., N.R., and F.L.

Investigation: G.D.M., V.P., F.A., A.V.P., S.K., L.K., A.C., M.T., A.P., and A.T. Funding Acquisition: G.D.M., M.L., P.M.L., N.R., F.L., and H.B.

Resources: B.S.T., D.H., N.W., S.M., R.K., E.S.L., N.R., V.T., and F.L. Supervision: H.B.

Writing – Original Draft: G.D.M., and H.B. Writing – Review & Editing: all authors

## REFERENCES

1. WHO. WHO Coronavirus Disease (COVID-19) Dashboard. https://covid19.who.int/ (2022).

2. Paterson, R.W., et al. The emerging spectrum of COVID-19 neurology: clinical, radiological and laboratory findings. Brain : a journal of neurology 143, 3104–3120 (2020).

3. Chou, S.H.-Y., et al. Global Incidence of Neurological Manifestations Among Patients Hospitalized With COVID-19—A Report for the GCS-NeuroCOVID Consortium and the ENERGY Consortium. JAMA Network Open 4, e2112131–e2112131 (2021).

4. Klimek, L., et al. Olfactory and gustatory disorders in COVID-19. Allergo Journal International (2022).

5. Thakur, K.T., et al. COVID-19 neuropathology at Columbia University Irving Medical Center/New York Presbyterian Hospital. Brain : a journal of neurology 144, 2696–2708 (2021).

6. Ruz-Caracuel, I., et al. Neuropathological findings in fatal COVID-19 and their associated neurological clinical manifestations. Pathology (2022).

7. Christensen, P.A., et al. Signals of Significantly Increased Vaccine Breakthrough, Decreased Hospitalization Rates, and Less Severe Disease in Patients with Coronavirus Disease 2019 Caused by the Omicron Variant of Severe Acute Respiratory Syndrome Coronavirus 2 in Houston, Texas. The American Journal of Pathology 192, 642–652 (2022).

8 Garrett, N., et al. High Rate of Asymptomatic Carriage Associated with Variant Strain Omicron. medRxiv, 2021.2012.2020.21268130 (2022).

9 Vihta, K.-D., et al. Omicron-associated changes in SARS-CoV-2 symptoms in the United Kingdom. medRxiv, 2022.2001.2018.22269082 (2022).

10. Chen, M., et al. Evolution of nasal and olfactory infection characteristics of SARS- CoV-2 variants. bioRxiv, 2022.2004.2012.487379 (2022).

11. Boscolo-Rizzo, P., et al. Coronavirus disease 2019 (COVID-19)–related smell and taste impairment with widespread diffusion of severe acute respiratory syndrome–coronavirus-2 (SARS-CoV-2) Omicron variant. International Forum of Allergy & Rhinology, 1–9 (2022).

12. Cardoso, C.C., et al. Olfactory Dysfunction in Patients With Mild COVID-19 During Gamma, Delta, and Omicron Waves in Rio de Janeiro, Brazil. JAMA (2022).

13. Butowt, R., Bilińska, K. & von Bartheld, C. Why Does the Omicron Variant Largely Spare Olfactory Function? Implications for the Pathogenesis of Anosmia in Coronavirus Disease 2019 The Journal of infectious diseases 226, 1304–1308 (2022).

14. de Melo, G.D., et al. COVID-19-related anosmia is associated with viral persistence and inflammation in human olfactory epithelium and brain infection in hamsters. Science Translational Medicine 13, eabf8396 (2021).

15. Chan, J.F., et al. Simulation of the clinical and pathological manifestations of Coronavirus Disease 2019 (COVID-19) in golden Syrian hamster model: implications for disease pathogenesis and transmissibility. Clin Infect Dis (2020).

16. Sia, S.F., et al. Pathogenesis and transmission of SARS-CoV-2 in golden hamsters. Nature 583, 834–838 (2020).

17. Käufer, C., et al. Microgliosis and neuronal proteinopathy in brain persist beyond viral clearance in SARS-CoV-2 hamster model. eBioMedicine 79, 103999 (2022).

18. Bauer, L., et al. Differences in neuroinflammation in the olfactory bulb between D614G, Delta and Omicron BA.1 SARS-CoV-2 variants in the hamster model. bioRxiv, 2022.2003.2024.485596 (2022).

19. Halfmann, P.J., et al. SARS-CoV-2 Omicron virus causes attenuated disease in mice and hamsters. Nature 603, 687–692 (2022).

20. Mohandas, S., et al. Pathogenicity of SARS-CoV-2 Omicron (R346K) variant in Syrian hamsters and its cross-neutralization with different variants of concern. eBioMedicine 79, 103997 (2022).

21. McMahan, K., et al. Reduced pathogenicity of the SARS-CoV-2 omicron variant in hamsters. Med 3, 262–268.e264 (2022).

22. Hayn, M., et al. Systematic functional analysis of SARS-CoV-2 proteins uncovers viral innate immune antagonists and remaining vulnerabilities. Cell reports 35, 109126 (2021).

23. Aliyari, S.R., et al. The Evolutionary Dance between Innate Host Antiviral Pathways and SARS-CoV-2. Pathogens 11, 538 (2022).

24. Su, C.-M., Wang, L. & Yoo, D. Activation of NF-κB and induction of proinflammatory cytokine expressions mediated by ORF7a protein of SARS-CoV-2. Sci Rep 11, 13464 (2021).

25. Fogeron, M.-L., et al. SARS-CoV-2 ORF7b: is a bat virus protein homologue a major cause of COVID-19 symptoms? bioRxiv, 2021.2002.2005.428650 (2021).

26. Kim, D.-K., et al. A map of binary SARS-CoV-2 protein interactions implicates host immune regulation and ubiquitination. bioRxiv, 2021.2003.2015.433877 (2021).

27. Mazur-Panasiuk, N., et al. Expansion of a SARS-CoV-2 Delta variant with an 872 nt deletion encompassing ORF7a, ORF7b and ORF8, Poland, July to August 2021. Euro Surveill 26, 2100902 (2021).

28. Panzera, Y., et al. A deletion in SARS-CoV-2 ORF7 identified in COVID-19 outbreak in Uruguay. Transboundary and emerging diseases 68, 3075–3082 (2021).

29. Jelley, L., et al. Genomic epidemiology of Delta SARS-CoV-2 during transition from elimination to suppression in Aotearoa New Zealand. Nature Commun 13, 4035 (2022).

30. Pyke, A.T., et al. Replication Kinetics of B.1.351 and B.1.1.7 SARS-CoV-2 Variants of Concern Including Assessment of a B.1.1.7 Mutant Carrying a Defective ORF7a Gene. Viruses 13(2021).

31. Renier, N., et al. iDISCO: A Simple, Rapid Method to Immunolabel Large Tissue Samples for Volume Imaging. Cell 159, 896–910 (2014).

32. Pepe, A., Pietropaoli, S., Vos, M., Barba-Spaeth, G. & Zurzolo, C. Tunneling nanotubes provide a route for SARS-CoV-2 spreading. Science Advances 8,

33. eabo0171 (2022).

33. Peyrin, J.-M., et al. Axon diodes for the reconstruction of oriented neuronal networks in microfluidic chambers. Lab on a Chip 11, 3663–3673 (2011).

34. Sainath, R. & Gallo, G. The dynein inhibitor Ciliobrevin D inhibits the bidirectional transport of organelles along sensory axons and impairs NGF-mediated regulation of growth cones and axon branches. Dev Neurobiol 75, 757–777 (2015).

35. Bauer, L., et al. The neuroinvasiveness, neurotropism, and neurovirulence of SARS- CoV-2. Trends in Neurosciences 45, 358–368 (2022).

36. Matschke, J., et al. Neuropathology of patients with COVID-19 in Germany: a post mortem case series. Lancet Neurol 19, 919–929 (2020).

37. Song, E., et al. Neuroinvasion of SARS-CoV-2 in human and mouse brainNeuroinvasion of SARS-CoV-2 in humans and mice. Journal of Experimental Medicine 218, e20202135 (2021).

38. Meinhardt, J., et al. Olfactory transmucosal SARS-CoV-2 invasion as port of Central Nervous System entry in COVID-19 patients. Nat Neurosci 24, 168–175 (2021).

39. Rutkai, I., et al. Neuropathology and virus in brain of SARS-CoV-2 infected non- human primates. Nature Commun 13, 1745 (2022).

40. Ferren, M., et al. Hamster organotypic modeling of SARS-CoV-2 lung and brainstem infection. Nature Commun 12, 5809 (2021).

41. Pellegrini, L., et al. SARS-CoV-2 Infects the Brain Choroid Plexus and Disrupts the Blood-CSF Barrier in Human Brain Organoids. Cell Stem Cell 27, 951–961.e955 (2020).

42. Ramani, A., et al. SARS-CoV-2 targets neurons of 3D human brain organoids. EMBO J 39, e106230 (2020).

43. Kong, W., et al. Neuropilin-1 Mediates SARS-CoV-2 Infection of Astrocytes in Brain Organoids, Inducing Inflammation Leading to Dysfunction and Death of Neurons. mBio 13, e0230822 (2022).

44. Khan, M., et al. Visualizing in deceased COVID-19 patients how SARS-CoV-2 attacks the respiratory and olfactory mucosae but spares the olfactory bulb. Cell (2021).

45. Helms, J., et al. Neurologic Features in Severe SARS-CoV-2 Infection. N Engl J Med 382, 2268–2270 (2020).

46. Yang, A.C., et al. Dysregulation of brain and choroid plexus cell types in severe COVID-19. Nature 595, 565–571 (2021).

47. Douaud, G., et al. SARS-CoV-2 is associated with changes in brain structure in UK Biobank. Nature 604, 697–707 (2022).

48. Abdelnabi, R., et al. The omicron (B.1.1.529) SARS-CoV-2 variant of concern does not readily infect Syrian hamsters. Antiviral Res 198, 105253 (2022).

49. Yuan, S., et al. Pathogenicity, transmissibility, and fitness of SARS-CoV-2 Omicron in Syrian hamsters. Science 0, eabn8939 (2022).

50. Hui, K.P.Y., et al. SARS-CoV-2 Omicron variant replication in human bronchus and lung ex vivo. Nature 603, 715–720 (2022).

51. Armando, F., et al. SARS-CoV-2 Omicron variant causes mild pathology in the upper and lower respiratory tract of hamsters. Nature Commun 13, 3519 (2022).

52. Reyna, R.A., et al. Recovery of anosmia in hamsters infected with SARS-CoV-2 is correlated with repair of the olfactory epithelium. Sci Rep 12, 628 (2022).

53. Urata, S., et al. Regeneration Profiles of Olfactory Epithelium after SARS-CoV-2 Infection in Golden Syrian Hamsters. ACS Chem Neurosci 12, 589–595 (2021).

54. Messlinger, K., Neuhuber, W. & May, A. Activation of the trigeminal system as a likely target of SARS-CoV-2 may contribute to anosmia in COVID-19. Cephalalgia 42, 176–180 (2022).

55. Kishimoto-Urata, M., et al. Prolonged and extended impacts of SARS-CoV-2 on the olfactory neurocircuit. Sci Rep 12, 5728 (2022).

56. Zazhytska, M., et al. Non-cell-autonomous disruption of nuclear architecture as a potential cause of COVID-19-induced anosmia. Cell 185, 1052–1064.e1012 (2022).

57. Bourgon, C., et al. Neutrophils initiate the destruction of the olfactory epithelium during SARS-CoV-2 infection in hamsters. bioRxiv, 2022.2003.2015.484439 (2022).

58. Schaecher, S.R., Touchette, E., Schriewer, J., Buller, R.M. & Pekosz, A. Severe Acute Respiratory Syndrome Coronavirus Gene 7 Products Contribute to Virus-Induced Apoptosis. Journal of Virology 81, 11054–11068 (2007).

59. Timilsina, U., Umthong, S., Ivey, E.B., Waxman, B. & Stavrou, S. SARS-CoV-2 ORF7a potently inhibits the antiviral effect of the host factor SERINC5. Nature Commun 13, 2935 (2022).

60. Schaecher, S.R., et al. An immunosuppressed Syrian golden hamster model for SARS-CoV infection. Virology 380, 312–321 (2008).

61. Silvas, J.A., et al. Contribution of SARS-CoV-2 Accessory Proteins to Viral Pathogenicity in K18 Human ACE2 Transgenic Mice. Journal of Virology 95, e00402–00421 (2021).

62. Imai, M., et al. Syrian hamsters as a small animal model for SARS-CoV-2 infection and countermeasure development. Proc Natl Acad Sci U S A 117, 16587–16595 (2020).

63. Bryche, B., et al. Massive transient damage of the olfactory epithelium associated with infection of sustentacular cells by SARS-CoV-2 in golden Syrian hamsters. Brain Behav Immun 89, 579–586 (2020).

64. Frere, J.J., et al. SARS-CoV-2 infection in hamsters and humans results in lasting and unique systemic perturbations post recovery. Science Translational Medicine 0, eabq3059 (2022).

65. Bulfamante, G., et al. Brainstem neuropathology in two cases of COVID-19: SARS-CoV-2 trafficking between brain and lung. Journal of Neurology 268, 4486–4491 (2021).

66. DosSantos, M.F., et al. Neuromechanisms of SARS-CoV-2: A Review. Frontiers in Neuroanatomy 14(2020).

67. Ueha, R., et al. Oral SARS-CoV-2 Inoculation Causes Nasal Viral Infection Leading to Olfactory Bulb Infection: An Experimental Study. Frontiers in cellular and infection microbiology 12(2022).

68. Dubé, M., et al. Axonal Transport Enables Neuron-to-Neuron Propagation of Human Coronavirus OC43. J Virol 92(2018).

69. Shi, J., et al. PHEV infection: A promising model of betacoronavirus-associated neurological and olfactory dysfunction. PLoS pathogens 18, e1010667 (2022).

70. Planas, D., et al. Reduced sensitivity of SARS-CoV-2 variant Delta to antibody neutralization. Nature 596, 276–280 (2021).

71. Zivcec, M., Safronetz, D., Haddock, E., Feldmann, H. & Ebihara, H. Validation of assays to monitor immune responses in the Syrian golden hamster (Mesocricetus auratus). J Immunol Methods 368, 24–35 (2011).

72. Gowen, B.B., et al. MP-12 virus containing the clone 13 deletion in the NSs gene prevents lethal disease when administered after Rift Valley fever virus infection in hamsters. Front Microbiol 6(2015).

73. Espitia, C.M., et al. Duplex real-time reverse transcriptase PCR to determine cytokine mRNA expression in a hamster model of New World cutaneous leishmaniasis. BMC Immunology 11, 31 (2010).

74. Planas, D., et al. Considerable escape of SARS-CoV-2 Omicron to antibody neutralization. Nature 602, 671–675 (2022).

75. Baer, A. & Kehn-Hall, K. Viral Concentration Determination Through Plaque Assays: Using Traditional and Novel Overlay Systems. JoVE, e52065 (2014).

76. Thi Nhu Thao, T., et al. Rapid reconstruction of SARS-CoV-2 using a synthetic genomics platform. Nature 582, 561–565 (2020).

77. Kouprina, N. & Larionov, V. Transformation-associated recombination (TAR) cloning for genomics studies and synthetic biology. Chromosoma 125, 621–632 (2016).

78. Kouprina, N., Noskov, V.N. & Larionov, V. Selective isolation of large segments from individual microbial genomes and environmental DNA samples using transformation-associated recombination cloning in yeast. Nature Protocols 15, 734–749 (2020).

79. Noskov, V.N., et al. A general cloning system to selectively isolate any eukaryotic or prokaryotic genomic region in yeast. BMC Genomics 4, 16 (2003).

80. Noskov, V.N., Lee, N.C.O., Larionov, V. & Kouprina, N. Rapid generation of long tandem DNA repeat arrays by homologous recombination in yeast to study their function in mammalian genomes. Biological Procedures Online 13, 8 (2011).

81. Lindenbach, B.D. Measuring HCV Infectivity Produced in Cell Culture and In Vivo. in Hepatitis C: Methods and Protocols (ed. Tang, H.) 329–336 (Humana Press, Totowa, NJ, 2009).

82. Lazarini, F., Gabellec, M.-M., Torquet, N. & Lledo, P.-M. Early Activation of Microglia Triggers Long-Lasting Impairment of Adult Neurogenesis in the Olfactory Bulb. The Journal of Neuroscience 32, 3652–3664 (2012).

83. AVMA. AVMA Guidelines for the Euthanasia of Animals: 2020 Edition*, (American Veterinary Medical Association, Schaumburg, IL, 2020).

84. Corman, V.M., et al. Detection of 2019 novel coronavirus (2019-nCoV) by real-time RT-PCR. Euro Surveill 25, 2000045 (2020).

85. Pfaffl, M.W. A new mathematical model for relative quantification in real-time RT-PCR. Nucleic Acids Res 29, e45–e45 (2001).

86. Renier, N., et al. Mapping of Brain Activity by Automated Volume Analysis of Immediate Early Genes. Cell 165, 1789–1802 (2016).

87. Gribaudo, S., et al. Propagation of alpha-Synuclein Strains within Human Reconstructed Neuronal Network. Stem Cell Reports 12, 230–244 (2019).

